# Disrupting USP39 deubiquitinase function impairs the survival and migration of multiple myeloma cells through ZEB1 degradation

**DOI:** 10.1101/2024.04.23.590548

**Authors:** Jessy Sirera, Saharnaz Sarlak, Manon Teisseire, Alexandrine Carminati, Coline Savy, Victoria J Nicolini, Patrick Brest, Thierry Juel, Christophe Bontoux, Marcel Deckert, Mickael Ohanna, Sandy Giuliano, Maeva Dufies, Gilles Pages, Frederic Luciano

## Abstract

**Rationale:** Multiple Myeloma (MM) stands as the second most common hematological malignancy characterized by the accumulation of monoclonal plasmocytes within the bone marrow. Despite the introduction of proteasome inhibitors, immunomodulatory agents and CD38-targeting antibodies which have extended survival rates, the disease remains incurable for most patients due to the emergence of resistant clones and frequent relapses. The efficacy of the proteasome inhibitor bortezomib (BTZ) in MM treatment underscores the critical role of the ubiquitin proteasome system (UPS) in this cancer. Deubiquitinases (DUBs), a class of enzymes governing the stability, interactions or localization of cellular proteins by removing ubiquitin modifications, have emerged as promising therapeutic targets across various cancers, including MM.

**Methods:** Through an exhaustive loss-of-function approach, we have identified for the first time USP39 DUB as a pivotal survival determinant for MM cells.

**Results:** Our analysis reveals a direct correlation between heightened USP39 mRNA levels and shorter survival in MM patients. Additionally, robust USP39 protein expression is observed in MM patient plasmocytes compared to healthy counterparts. Knockdown of Usp39 not only impedes clonogenic capabilities, but also induces apoptosis, triggers cell cycle arrest and overcomes BTZ resistance. Complementary gain-of-function assays, further elucidate how USP39, by stabilizing the transcription factor ZEB1, enhances the trans-migratory potential of MM cells.

**Conclusions:** In summary, our findings underscores the pivotal role of the deubiquitinase USP39, suggesting that targeting the USP39/ZEB1 axis hold promise as a prospective diagnostic marker and therapeutic target in MM.

## INTRODUCTION

Multiple myeloma (MM) is a rare blood disease, comprising 10% of all hematological malignancies, and ranks as the second most prevalent blood cancer [1]. While MM has traditionally been associated with the elderly, with the majority of patients diagnosed between the age of 60 and 70, there has been a noticeable increase in diagnoses among younger individuals in recent years. The management of (MM) has advanced significantly over the past decade with the approval of novel therapeutic agents such as proteasome inhibitors (PIs), immunomodulatory drugs (IMiDs), monoclonal antibodies (mAbs) and CAR-T-cell therapy [2]. Refinements in treatment combinations have further prolonged the overall survival of MM patients, with some experiencing survival times of up to 7 years or more. Despite these advancements, the emergence of resistance to one or more of these treatment modalities often leads to relapses or the development of refractory disease [3]. Consequently, there is a pressing need to identify new therapeutic targets that play pivotal roles in the initiation and progression of MM.

The ubiquitin-proteasome system (UPS) is instrumental in maintaining cellular homeostasis by facilitating the degradation of a majority of short-lived proteins, thereby eliminating misfolded, damaged, and potentially harmful proteins [4]. This system plays a crucial role in various biological processes including cell signaling, receptor trafficking, cell cycle regulation, DNA damage, cell proliferation, selective autophagy and apoptosis by selectively targeting cellular proteins for degradation [5]. Dysfunctions in the UPS have been implicated in numerous diseases, including hematological malignancies [6]. The use of proteasome inhibitors (e.g., BTZ or carfilzomib) to inhibit UPS activity has a key therapeutic strategy in MM [7]. Malignant plasma cells in MM patients are particularly sensitive to proteasome inhibition due to their high production of misfolded and non-functional proteins, which must be degraded by the proteasome to prevent the formation of toxic protein aggregates [8]. The UPS comprises essential components such as ubiquitinases (Ubis), deubiquitinases (DUBs) and the 26S proteasome. Ubiquitination involves the covalent attachment of ubiquitin molecules (Ub) to a lysine moieties residues to target proteins. This process is mediated by activating (E1), conjugating (E2), and ligase (E3) enzymes resulting in mono- or polyubiquitination on seven lysine residues (K6, K11, K27, K29, K33, K48, and K63) of target protein. K48 polyubiquitinated chains have historically been identified as primarily responsible for proteasome degradation of short-lived proteins and ubiquitin recycling [9].

DUB play a crucial role in regulating ubiquitination dynamics by removing of ubiquitin moieties from substrates thereby modulating protein stability, interaction and function [10]. DUBs encompass approximately 100 family proteins classified into six families based on their domain structure including ubiquitin-specific proteases (USPs), ubiquitin carboxy-terminal hydrolases (UCHs), Machado-Joseph disease protein domain proteases (MJDs), ovarian tumor-related proteases (OTUs), and JAB1/PAB1/MPN-domain containing metallo-enzyme (JAMMs) [11].

Emerging evidence suggests that dysregulation of DUBs contributes significantly to cancer pathogenesis, particularly in hematological malignancies [12]. Elevated level of specific DUBs are often observed in cancers leading to the stabilization of pro-tumor factors. In MM, DUBs such as USP5 and Otub1 stabilize the oncogene c-Maf promoting myeloma cell proliferation and survival [13], [14]. Additionally, USP9X stabilizes the anti-apoptotic protein Mcl-1, enhancing cell survival [15] while USP7 contributes to BTZ resistance by stabilizing NEK2 protein [16].

Ubiquitin-specific peptidase 39 (USP39), a member of the DUBs family, possesses both the UBP-type zinc finger and ubiquitin C terminal hydrolase domains [17]. Moreover, aberrant USP39 expression is linked to tumorigenesis in various cancers including breast [18], colorectal [19], lung [20], and brain [21] cancers. Six substrates are regulated by USP39 including CHK2 kinase, SP1 transcription factor, Stat1, FOXM1 transcription factor, cyclin B1 and the Zinc-finger E-box-binding homeobox 1 (ZEB1), which is a pivotal inducer of epithelial-to-mesenchymal transition (EMT). Their stabilization by USP39 induces chemo-radiation resistance [22], promotes tumorigenesis in hepatocellular carcinoma (HCC) [17], sustains IFN-Induced antiviral immunity [23], facilitates breast cancer cell proliferation [24], promotes tumor cell proliferation and glioma and HCC progression [25] [26], respectively.

In our study, employing a comprehensive loss-of-function approach, we identified USP39 as a critical survival factor for MM cells. Elevated USP39 mRNA levels correlated with shorter survival of MM patients and heightened USP39 protein levels were observed in MM patients plasmocytes compared to healthy individuals. Down-regulation of Usp39 inhibited clonogenic abilities, induced apoptosis, cell cycle arrest and overcame BTZ resistance in MM cells. Moreover, gain-of-function experiments revealed that USP39 promoted MM cells trans-migratory capacity through ZEB1 stabilization. Our findings underscore the significance of USP39 in MM pathogenesis and suggest that targeting the USP39/ZEB1 axis hold promise as a therapeutic strategy in MM.

## MATERIAL AND METHODS

### Cell lines

The OPM2, U266 (TIB-196), RPMI8226 (CCL-155) and MM1S (CRL-2974) and AF-10 human MM cell lines were purchased from the ATCC. The LP1 (ACC 41) MM cell line was obtained from the DSMZ. All cell lines were grown at 37°C under 5% CO2 in RPMI medium (Gibco BRL, Paisley, UK) supplemented with 10% fetal calf serum (Gibco BRL, 10270), 50 units/ml penicillin, 50 mg/ml streptomycin (Lonza, DE17-602E) and 1 mM sodium pyruvate (Lonza, BE13-115E). U266R cells resistant to BTZ were previously described [27] and were maintained in culture under similar conditions as the parental U266 cell line. HS-5 stromal cells (CRL-6311) were purchased from the ATCC and cultured at 37°C under 5% CO2 in DMEM medium (Gibco BRL, Paisley, UK) supplemented with 10% fetal calf serum (Gibco BRL, 10270), 50 units/ml penicillin, 50 mg/ml streptomycin (Lonza, DE17-602E).

### Reagents

Sodium fluoride (S7920), sodium orthovanadate (220590), phenylmethylsulfonyl fluoride (PMSF) (P7626), aprotinin (A162-B), leupeptin (SP-04-2217-B), Triton X-100 (N150) was purchased from Sigma. MG132 (BMLPI102) and CHX (ALX-380-269-G001) were purchased from Enzo Life Sciences. Bortezomib (CAS 179324-69-7) was purchased from Calbiochem.

### Antibodies

Anti-PARP (#9542), anti-Chk2 (#6334), anti-Stat1 (#9172), anti-Myc-Tag (#2276), anti-β-catenin (#8480), anti-N-cadherin (#4061), anti-vimentin (5741), and anti-rabbit HRP conjugated (#7074) antibodies were purchased from Cell Signaling Technology. Anti-USP39 antibody (A304-817A-T) used for immunoblotting was purchased from Bethyl Laboratories. Anti-USP39 (ab245571), and (ab131244) antibodies used for immunoprecipitation and IHC experiments respectively were purchased from Abcam. GAPDH (ab9485) and SP1 (ab227383) were purchased from Abcam. Anti-ZEB1 antibody (21544-1-AP) was purchased from Proteintech. Anti-HA antibody (sc-53516) was purchased from Santa Cruz Biotechnology).

### Immunoblotting

After stimulation, the cells were lysed at 4°C in lysis buffer (50 mM HEPES pH 7.4, 150 mM NaCl, 20 mM EDTA, 100 μM NaF, 10 μM Na3VO4, 1 mM PMSF, 10 μg/ml leupeptin, 10 μg/ml aprotinin, and 1% Triton X-100). The lysates were centrifuged at 16,000 ×g for 15 min at 4°C, and the supernatants were supplemented with concentrated SDS sample buffer. A total of 30 μg of protein was separated on a 12% polyacrylamide gel and transferred onto a PVDF membrane (Immobilon-P, Millipore, IPVH00010) in a 20 mM Tris, 150 mM glycine and 20% ethanol buffer at 250mA for 1h30 min at 4°C. After blocking the non-specific binding sites in saturation buffer (50 mM Tris pH 7.5, 50 mM NaCl, 0.15% Tween, and 5% BSA), the membranes were incubated with the specific antibodies, washed three times using TNA-1% NP-40 (50 mM Tris pH 7.5, and 150 mM NaCl) and incubated further with HRP-conjugated antibody for 1h at room temperature. The immunoblots were revealed using the enhanced chemiluminescence detection kit (Pierce, 32106).

### Plasmids

USP39 Myc-DDK plasmid (NM_006590) Human Tagged ORF Clone (RC209551) was purchased from OriGene. HA-ubiquitin plasmid (#18712) was purchased from Addgene. P3-Flag-CMV-10-ZEB1 (PVTB00266-2b) plasmid was purchased from Bioactiva Diagnostica.

Construction of the pPRIPu USP39-MYC. The pPRIPu CrUCCI backbone (kind gift from Dr. F Delaunay) was amplified with primer adaptors for AgeI and BamH1. USP39-MYC was amplified by PCR with primer adaptors for AgeI and BglII (sticky end compatible with BamHI) from USP39 Myc-DDK (RC209551) plasmid. The USP39-MYC fragment was inserted in pPRIPU retroviral vector. The entire sequence in the latter has been confirmed by sequencing analysis. Briefly, replication-defective, self-inactivating retroviral constructs were used to establish stable OPM2 and U266 cell lines. Selection was performed by puromycin (4 µg/ml). Then, the cells were sorted as a polyclonal population and used in the following experiments.

### Plasmid transfection

Briefly, 3 million KMM1 cells were transfected with 2 μg of corresponding plasmids using JetPEI reagent (Polyplus, 101000053). Then, the cells were plated in 3ml of RPMI 10% FCS media and incubated at 37°C until experiment analysis.

### siRNAs

USP39 small interfering RNAs (siRNA) were purchased from Invitrogen life technologies: USP39#1 (GGGUAUUGUGGGACUGAAUAACAUA); USP39#2 (CCAGACAACUAUGAGAUCAUCGAUU); USP39#3 (UCAUGUUCUUGUUGGUCCAGCGUUU). Zeb1 siRNAs were purchased from Dharmacon (ON-TARGET plus SMART pool siRNA J-006564). Transfection of U266 cells was performed as described previously [49] using the Nucleofector system (Lonza, VCA-1003). Briefly, 2.5 million cells were electroporated with either control or HSPB8 siRNA (100nM) using nucleofector (kit C and program X-05). Then, the cells were plated in 5 ml of RPMI 10% FCS media and incubated for 48h at 37°C until experiment analysis. Transfections of KMM1 and OPM2 cells were performed using Lipofectamine RNAimax reagent (Invitrogen; 13778075) in accordance with the manufacturer’s instructions.

The Human ON-TARGETplus siRNA Library - Deubiquitinating Enzymes (G-104705) was purchased from Horizon Discovery.

### SiRNAs transfection

2.5 million of U266 cells were electroporated with either control, USP39 or ZEB1 siRNA (100 nM) using nucleofector (kit C and program X-05). Then, the cells were plated in 5 ml of RPMI 10% FCS media and incubated for 48h at 37°C until experiment analysis.

Transfections of KMM1 and OPM2 cells were performed using Lipofectamine RNAimax reagent (Invitrogen; 13778075) in accordance with the manufacturer’s instructions.

### Production and infection of Myc-USP39 retroviral particles

Replication-defective, self-inactivating retroviral constructs were used for establishing stable OPM2 and U266 cell lines. On day 1, HEK293T were seeded in T25 flask (500 000 cells). On day 2, cells were co-transfected with 3,33μg transfer (pPRIPu USP39-MYC), 1,66μg packaging (pCMV-gag-pol) and 1,66 μg envelope (pCMV-env-VSV-G) plasmids, using lipofectamine 3000. After 16 h, medium was replaced with 5ml fresh medium (day 3). Supernatant was harvested 48h after transfection and filtered on 0.45 μm PES filters to remove cell debris. 5 million of OPM2 and U266 cells were seeded and directly infected by applying filtered supernatant + 4μg/ml polybrene (Sigma-Aldrich) to the cells (day 4). Viruses were left for 24h before adding fresh media for 2 days. Then, infected cells were splitted and incubated with puromycine (1 μg/ml) for selection. Cells were frozen 20 days later or used in subsequent experiments.

### Infection of GFP-USP39 lentiviral particles

USP39 (NM 006590) human mGFP-tagged ORF clone lentiviral particles (RC209551L4V) and control lentiviral particles (PS100093V) were purchased from OriGene.

5 million of OPM2 and U266 cells were seeded and directly infected by applying filtered supernatant + 4 μg/ml polybrene (Sigma-Aldrich) to the cells (day 4). Viruses were left for 3 days before splitting infected cells and adding puromycin (1 μg/ml) for selection. Cells were frozen 20 days later or used in subsequent experiments.

### Transmigration

The transmigration assay based on chemotaxis was performed by inserting a transwell polyester membrane filter with 8 μm pores polycarbonate membrane (Falcon® Cell Culture Inserts. Product number: 353097) in 24-well culture plates. Total of 150 000 serum starved U266, OPM2 and KMM1 cells were resuspended in 300 μL RPMI seeded in upper chamber of Transwell. To chemoattract KMM1 and OPM2 cells, RPMI medium containing 10% FBS was placed in the lower chamber. For U266 cells, 100 μg/mL of recombinant human stromal cell-derived factor 1alpha (SDF-1α) (PeproTech) was added. After 24h of incubation, the migrated cells on the bottom side of the membrane were counted. Each assay was performed in triplicate wells.

### ZEB1 Deubiquitination assay

KMM1 cells were transfected for 48h with HA-ub plasmid in the presence or in the absence of Myc-USP39 plasmid. Then cells were treated with MG132 at 1 µM for 8h. Cells were lysed in deubiquitination buffer (50 mM Tris-HCl pH 8.0, 50 mM NaCl, 1 mM EDTA, 10 mM DTT, 5% glycerol, and fresh proteinase inhibitors). Then ubiquitinated ZEB1 was purified from cell extracts with non relevant IgG or anti-ZEB1 antibody. Inputs were immunoblotted with anti-HA and anti-Myc antibodies to visualize poly-HA-Ub and USP39 respectively. IPs products were immunoblotted with anti-HA and anti-ZEB1 antibodies to visualize Ub-ZEB1 complex and immunoprecipitated ZEB1.

### IncuCyte screening

The Human ON-TARGET plus siRNA Library - Deubiquitinating Enzymes (Horizon Discovery, G-104705) was used to individually deplete each DUB on OPM2 cells. Briefly, 100 000 OPM2 cells were transfected with individual DUB siRNA (20 nM) using Lipofectamine RNAimax reagent and seeded in 2.5 ml of RPMI medium without antibiotic in a 24 well plates. Cell confluence and cell death were monitored over a period of 5 days. Cell proliferation was determined as percent confluence from phase images and was analyzed by IncuCyte image analysis software (Sartorius). For cell death assays, PI (10 μg/ml) was added to the medium, and cell death was calculated as red objects normalized to the confluency factor and the initial timepoint. Mcl-1 siRNA was used as a positive control of siRNA transfection efficiency as well as a normalization element for the different experiments. BTZ was used as cell death inducer.

### Colony formation assay

Cells were transfected with siRNAs or stimulated with BTZ (50 nM). The day after, 5000 cells were resuspended within a mix of methylcellulose (MethoCult™ H4100) and RPMI medium supplemented with L-Glutamine 1% and SVF 5% without antibiotics. Then, 500 μL of mix were placed in wells of a 24-well plate. Clones were stained with MTT reagent after 10 days. The number and size of the clones were analyzed using Image J software.

### IHC

Immunohistochemistry for USP39 (Anti-USP39 antibody ab131244) were performed using Myeloma tissue array (BM291d and BM483b) purchased from TissueArray.com. Antigen retrieval was performed by boiling sections for 10 min in citrate buffer (pH 6.0) and cooling at RT°, followed by blocking of endogenous peroxidase activity with 0.3% H_2_O_2_ for 30 min. The sections were blocked with 2.5% horse serum in TBS solution for 30 min in a humid chamber prior to incubation with anti-USP39 antibody (1/200). Positive cells were detected using an ImmPRESS HRP anti-rabbit detection kit. The immune complexes were visualized using a Peroxidase Substrate DAB kit (Vector) according to the manufacturer’s protocol, and slides were counterstained with hematoxylin. Blind quantification of brown staining was done as follows: Nuclear staining intensity were interpreted by pathologist visual scoring as - = no staining, + = low intensities, ++ = medium intensities, and +++ = high intensities. The % of USP39 positive cells was determined by ImageJ quantification and confirmed by pathologist visual scoring.

### Cell cycle

Cells were harvested, washed, and resuspended in cold 70% ethanol overnight. After two washes with PBS, cells were resuspended in propidium iodide 1,25 µg/mL (Biolegend, 421301) containing ribonuclease A (25 µg/mL) (Sigma, R4642) for fifteen minutes at room temperature and were analyzed using MACSQUANT Analyser (Myltenyi Biotech, 130-092).

### Co-immunoprecipitations

OPM2 cells were suspended in lysis buffer [50 mM TRIS-HCl, pH 7.4, 150 mM NaCl, 20 mM EDTA, 50 μM NaF, 0.5% NP-40, 10 μM Na3VO4, 20 µg ml−1 leupeptin, 20 µg ml−1 aprotinin, 1 mM dithiothreitol and 1 mM phenylmethyl sulfonyl fluoride (PMSF)]. The lysates (500 µl) were then incubated with either 1 μg of non relevant, anti-USP39 (ab245571) or anti-ZEB1 (21544-1-AP) antibodies and 30 µl protein G Sepharose (Zymed, 10-1242) at 4°C overnight. The beads were washed five times with 1 ml lysis buffer before boiling in Laemmli sample buffer. 30 µg of total lysates and immunoprecipitates were analyzed by immunoblotting using anti-USP39 (A304-817A-T) or ZEB1 (21544-1-AP) antibodies.

### Measurement of cell metabolism (XTT)

MM cells lines were incubated in a 96-well plate and then subjected to different experimental conditions. 50μl of the XTT reagent (Roche Applied Science, 11-465-015) (sodium 3’-[1-(phenylaminocarbonyl)-3,4-tetrazolium]-bis(4methoxy-6-nitro) benzene sulfonic acid hydrate) was added to each well. The assay is based on the cleavage of the yellow tetrazolium salt XTT to form an orange formazan dye by metabolically active cells. The absorbance of the formazan product, reflecting cell viability, was measured at 490 nm. Each assay was performed in triplicate.

### Cell Death assay

Cell viability was measured using the propidium iodide (PI) dyed exclusion assay. Briefly, after treatment, the cells were collected and incubated with PI (10 μg/ ml) for 5 min. The percentage of PI positive cells was next analyzed by flow cytometry using MACSQUANT Analyser (Miltenyi Biotech, 130-092).

### Caspase activity

Following treatments, cells were lysed for 30 min at 4°C in lysis buffer (50 mM HEPES pH 7.4, 150 mM NaCl, 20 mM EDTA, 1 mM PMSF, 10 μg/ml leupeptin, 10 μg/ml aprotinin and 0.2% Triton X-100), and lysates were cleared at 16,000 ×g for 15 min at 4°C. Each assay (in triplicate) was performed with 50 μg of protein prepared from control or stimulated cells. Briefly, cellular extracts were then incubated in a 96-well plate, with 0.2 mM of Ac-DEVD-AMC as substrates for various times at 37°C as previously described. Caspase activity was measured by following emission at 460 nm (excitation at 390 nm) in the presence or in absence of 10 μM of Ac-DEVD-CHO. Each experiment was performed in triplicates and repeated at least 3 times.

### Zebrafish tumor model

Zebrafish embryos of the transgenic strain expressing enhanced GFP under the fli1 promoter (Fli1:EGFP) were cultivated at a temperature of 28°C under standard experimental conditions. At 24h post fertilization (hpf), zebrafish embryos were exposed to an aquarium solution containing 1X embryo medium (5 mM NaCl, 0.16 mM KCl, 0.4 mM CaCl2, 0.4 mM MgSO4) with methylene blue solution. Upon reaching 48 hpf, Fli1:EGFP zebrafish embryos underwent dechorionation using sharp-tip forceps and were anesthetized with 0.04 mg/mL of tricaine (MS-222, Sigma) prior to microinjection. In vitro, OPM-2 cells were labelled with 2 μg/mL of Vybrant DiD cell-labeling solution (Life Technologies). The labeled cells were suspended in RPMI containing 2 mM EDTA. Subsequently, 5 nl of the cell solution was injected into the perivitelline space (PVS) of each embryo utilizing an Eppendorf microinjector (Femto-Jet 5247, Eppendorf) and a Manipulator MM33-Right (Märzhäuser Wetziar). Non-filamentous borosilicate glass capillary needles were employed for the injection procedure. The injected zebrafish embryos were promptly transferred to 1X embryo medium with methylene blue solution. Over the course of 48h, fluorescent microscopy (EVOS M5000) was used to monitor zebrafish embryos, investigating tumor invasion and metastasis.

### Statistical analysis

Results are expressed as mean ± SD and analyzed using GraphPad Prism 9 (RRID:SCR_002798) software. The unpaired Student’s *t* test was used to determine the difference between 2 independent groups. For multiple comparison analyses, one-way ANOVA test with Turkey’s correction was used. Evaluation of the gaussian distribution of the data was performed prior to the *t* test or ANOVA. Normal distribution of the data and variance similarity were verified using GraphPad Prism. A *P* value less than 0.05 was considered statistically significant. All experiments were performed with a minimum of *n* = 3 biological replicates and *n* = 3 technical replicates. For *in vivo* studies, two-tailed Mann-Whitney U test was utilized to compare two independent groups and the sample size was determined using the methods described by Berndtson et al [28]. Predetermined exclusion criteria included the absence of signal at the start of the experiment.

## RESULTS

### The depletion of USP39 suppresses proliferation and induces cell death in OPM2 myeloma cell line

To screen critical DUBs involved in MM proliferation, we utilized a DUB siRNA library (Dharmacon Company, Cat: G104705) containing 100 siRNAs specific for each DUB. Four siRNAs targeting a single DUB were transfected into the MM OPM2 cell line, and the percentage of cell confluence was monitored over time using Incucyte device (**Figure 1A**). The graph illustrates the proliferation index of OPM2 cells subjected to DUBs siRNA transfection, with a value of 1 value representing the proliferation rate of OPM2 cells transfected with control siRNA transfection. Values above 1 indicate an increase in proliferation rate following DUBs siRNA transfection, while values under 1 signify a decrease in cell proliferation rate. Our focus was on DUBs that reduced proliferation when inhibited (red histograms). Among them, several reported DUBs were identified in our screening assay, including PSMD14 and USP10. Notably, USP39 emerged as the top DUB significantly suppressing cell proliferation when inhibited (**Figure 1B**). Phase-contrast images revealed that siUSP39 led to both reduced OPM2 cell confluence and cell shrinkage, reminiscent of the effects observed following stimulation with the proteasome inhibitor BTZ (**Figure 1C**). By quantifying confluence (**Figure 1D**) and the percentage of dead cells (**Figure 1E**), we showed that siUSP39 decreased confluence and induced cell death starting from 48h. The graph illustrates the mean slope of OPM2 cell confluence following siControl, siUSP39, or BTZ (50nM) stimulation (**Figure 1F**). In summary, through this DUB loss-of-function screening, we identified USP39 as a critical factor essential for the proliferation and viability of MM cells.

**Figure 1:**
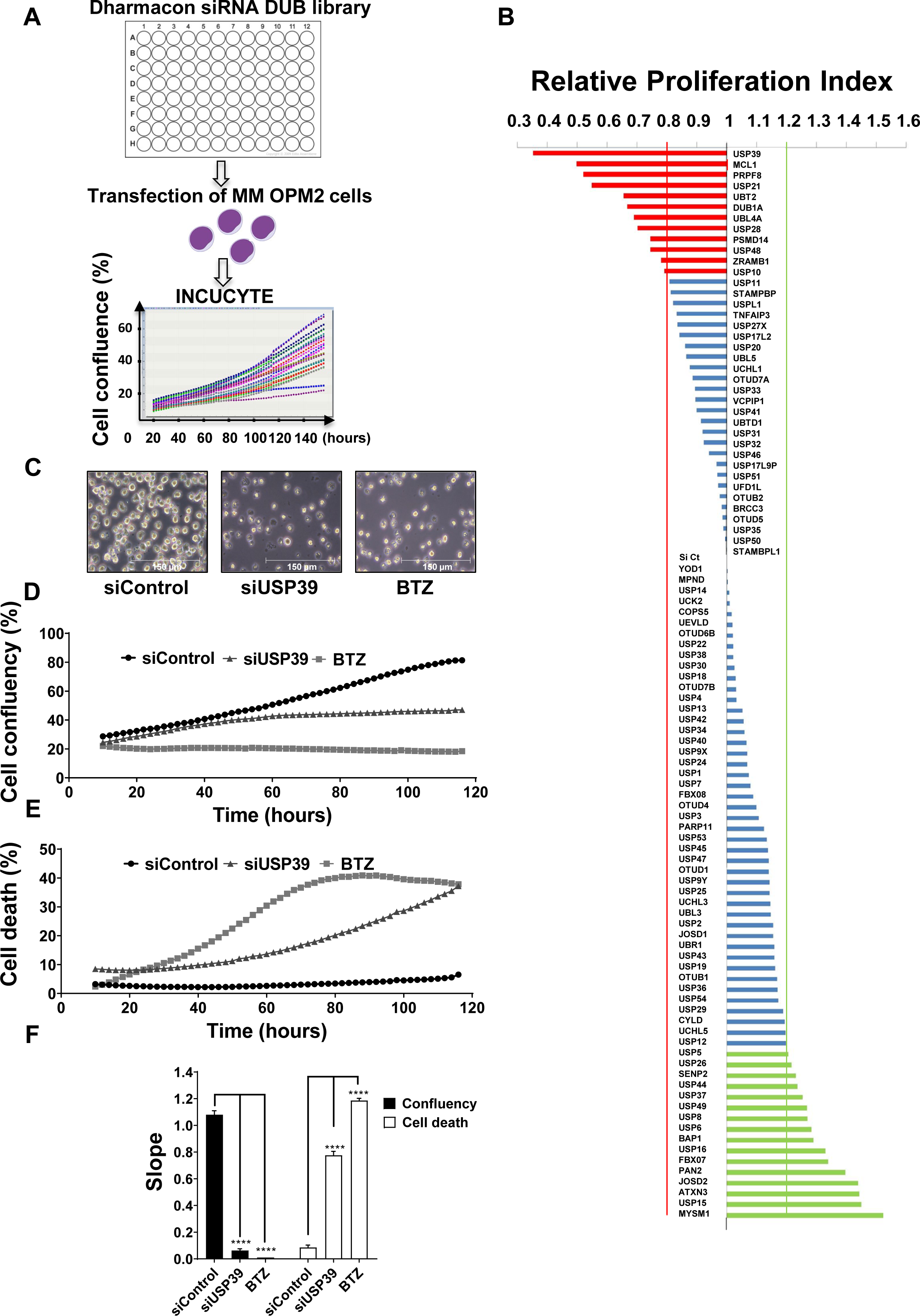
The depletion of USP39 suppresses proliferation and induces cell death in OPM2 myeloma cell line. **(A)** Schematic representation of siRNA screen using Dharmacon siRNA DUB library. Three siRNAs targeting a single DUB were transfected per well in MM OPM2 cell line and the percentage of cell confluence was measured over time using the Incucyte device. **(B)** Graph representing the proliferation index of OPM2 cells subjected to DUBs siRNA transfection. A value of 1 represents the proliferation rate of OPM2 subjected to Control siRNA transfection, while values above 1 indicate an increase in proliferation rate following DUBs siRNA transfection. Values below 1 indicate a decrease in cell proliferation rate following DUBs siRNA transfection. **(C)** Representative phase-contrast images of OPM2 cells transfected with Control or USP39 siRNAs for 96h or stimulated by BTZ (50nM) for 48 hours. **(D,E)** OPM2 cells were transfected with Control or USP39 siRNA, or stimulated by BTZ (50nM). The percentage of cell confluency **(D)** and the percentage of cell death were measured from 12 hours to 120 hours using the Incucyte device. **(F)** Graph representing the mean slope of OPM2 cell confluence following either siControl, siUSP39, or BTZ stimulation (50 nM). The results are presented as the mean of at least three independent experiments ± standard deviation (SD). Statistical analysis was performed using one-way ANOVA to compare differences between the sicontrol and the other two groups (siUSP39 or BTZ). Statistical significance is denoted as follows: *P < 0.05, **P < 0.01, ***P < 0.001, ****P < 0.0001, NS (non-significant).

### USP39 is over-expressed in MM patients compared to healthy donors and its high expression is correlated with shorter survival

Survival analysis conducted through Gene Set Enrichment Analysis (GSEA) revealed a significant correlation between high expression of USP39 mRNA and shorter survival among patients with MM (**Figure 2A**). Moreover, higher USP39 mRNA levels were observed in patients across various stages of the disease compared to healthy individuals (**Figure 2B**). To evaluate USP39 expression at the protein level, we conducted immunohistochemistry (IHC) experiments using bone marrow (BM) samples from MM patients and healthy individuals. Confirmation of BM samples from MM patients containing more CD138 staining in IHC experiments, whereas healthy individuals exhibited less than 2% plasma cells, as expected **(Figure S1)**. Notably, USP39 protein expression was primarily detected at the nuclear level in MM patients, while no expression was observed in healthy individuals (**Figure 2C**). Expanding our analysis with a larger cohort of MM BM samples, and in concordance with pathologists, we categorized the intensity of USP39 expression into four levels (+++ strong; ++ moderate; + weak; - undetectable) (**Figure 2D**). Among 12 MM patients examined, USP39 expression was detected in 9 individual, with staining intensity ranging from weak to strong, and the percentage of positive cells ranging from 46 to 92%. However, USP39 expression was not detected in 3 MM patients, nor in any healthy individuals (**Figure 2D**). Collectively, these findings indicate that both USP39 mRNA and protein are prominently expressed in MM patients compared to healthy individuals, with high expression correlating with shorter survival times.

**Figure 2:**
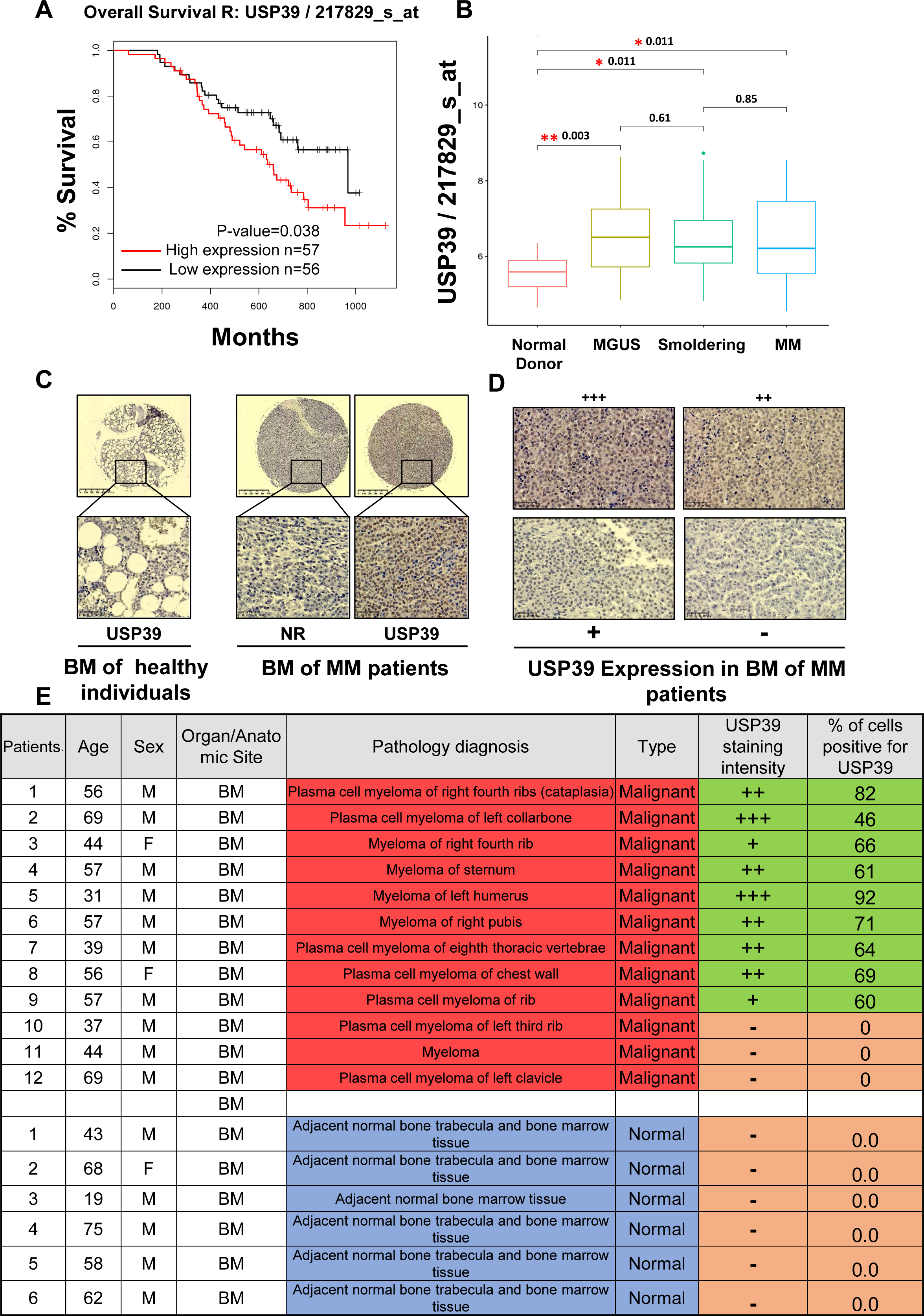
USP39 is overexpressed in MM patients compared to healthy donors and its high expression is correlated with shorter survival. **(A)** Kaplan-Meier of overall survival in patients with MM with high (red line) or low (black line) USP39 mRNA expression (P=0.038) (GSE9782). **(B)** USP39 mRNA expression in normal donor, MGUS, Smoldering and MM patients (GSE6477). **(C) Left**, Representative USP39 staining of bone marrow samples from healthy individuals **Right**, Representative USP39 staining of bone marrow samples from MM patients. “NR” denotes a non-relevant antibody. **(D)** Representative USP39 immunostaining of bone marrow samples from MM patients. Staining intensity was interpreted by a pathologist using visual scoring: “–” denotes undetectable, “+” denotes low intensities, “++” denotes medium intensities, and “+++” denotes high intensities. **(E)** Table representing USP39 staining of 12 bone marrows from MM patients and 6 bone marrows from healthy individuals. The percentage of USP39 positive cells was determined by ImageJ quantification and confirmed by pathologist visual scoring. Staining intensity were interpreted by pathologist visual scoring: “–“ denotes undetectable, “+” denotes low intensities, “++” denotes medium intensities, and “+++” denotes high intensities. The result is presented as the mean of each group of donors ± SD. Statistical analysis was performed using one-way ANOVA to compare differences between the Normal donor, MGUS, Smoldering and MM patients. Statistical significance is denoted as follows: *P < 0.05, **P < 0.01, ***P < 0.001, ****P < 0.0001, NS (non-significant).

### USP39 Depletion Suppresses Cell Proliferation, Induces Apoptosis, and decreases Clonogenicity in OPM2 and KMM1 Multiple Myeloma Cells

Prior to delving into the specifics of the USP39 targeting study, an assessment of USP39 mRNA expression was conducted using publicly available databases (DepMap Data Explorer), revealing high expression levels of USP39 mRNA across all human MM cell lines **(Figure S2A)**. These findings were further validated at the protein level through immunoblot analysis of USP39 in a selected panel of human MM cell lines **(Figure S2B)**.

To corroborate the results obtained from the loss-of-function screening, the study was extended to include a second MM cell line (KMM1) and two additional pairs of USP39 siRNAs. Specific inhibition of USP39 by these siRNAs led to a significant reduction in USP39 expression, accompanied by a substantial induction of cell death and a marked decrease in cellular metabolism in both OPM2 and KMM1 cells after 96 hours (**Figure 3A-B**). Furthermore, inhibition of USP39 by siRNA resulted in a diminished clonogenic capacity in both cell lines, with the proteasome inhibitor BTZ serving as a positive control for inhibition of clonogenic potential (**Figure 3C-D**). To confirm the specificity of the observed effects, a complementation experiment was conducted in KMM1 cells. Consistent with expectations, siRNA-mediated knockdown of endogenous USP39 resulted in a decrease in cellular metabolism after 96 hours. Transfection of a plasmid encoding an exogenous form of USP39 (Myc-USP39) alone did not affect cellular metabolism. However, co-transfection of USP39 siRNA and Myc-USP39 plasmid led to a partial rescue of the decline in cellular metabolism, thus demonstrating the specificity of the observed effects to USP39 modulation (**Figure 3E**). In summary, our findings indicate that targeted inhibition of USP39 results in decreased cellular metabolism, clonogenic potential, and induces late-stage cell death in various MM cell lines.

**Figure 3:**
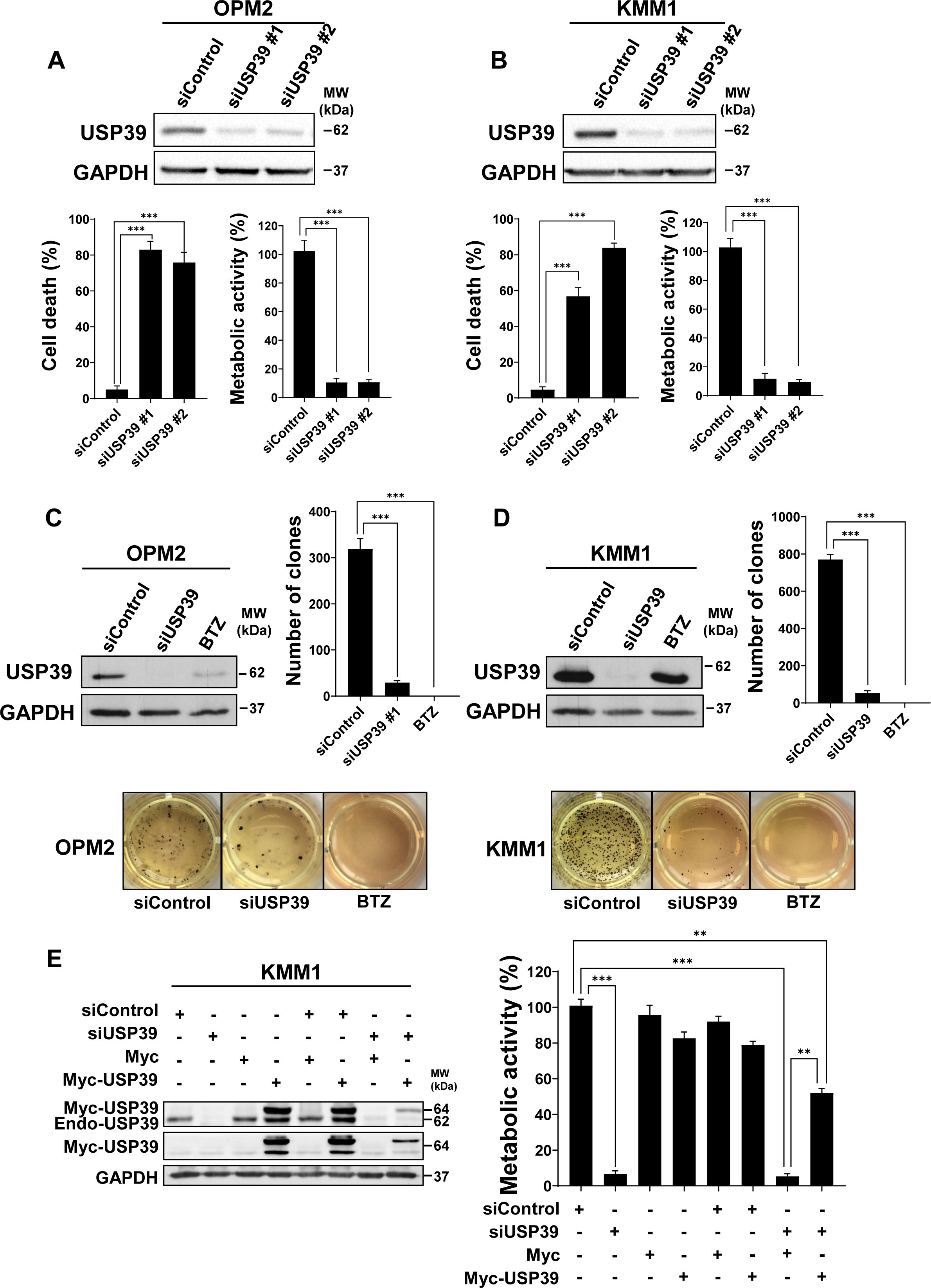
USP39 Depletion Suppresses Cell Proliferation, Induces Apoptosis, and decreases Clonogenicity in OPM2 and KMM1 Multiple Myeloma Cells. **(A)** OPM2 cells were transfected with either control or two different single USP39 siRNA (siUSP39 #1 and siUSP39 #2) for 96h. Then, lysates from these cells were subjected to immunoblots using GAPDH and USP39 antibodies (upper part). In parallel, the percentage of cell death was measured by flow cytometry after IP staining (left lower part) and cell metabolism was assessed by XTT assay (right lower part). **(B)** KMM1 were treated as described for OPM2 cells and subjected to the same analysis. **(C)** OPM2 cells were transfected with either control or single USP39 siRNAs for 96 hours or stimulated with BTZ for 48 hours. Lysates from these cells were subjected to immunoblots using GAPDH and USP39 antibodies (upper left). In parallel, the clonogenic capacity of the cells was measured after 10 days within a semi-solid medium. The quantification of the clonogenic assay is reported in the upper right part of the figure. Representative pictures were shown in the lower part. **(D)** KMM1 were treated as described for OPM2 cells and subjected to the same analysis. **(E)** KMM1 cells were either transfected with Control or USP39 siRNAs for 24 hours. Then cells were transfected with Myc-Tag or Myc-USP39 vectors. After 72 hours, lysates from these cells were subjected to immunoblots using GAPDH, myc-Tag or USP39 antibodies (left part). After 96 hours of transfections, cell metabolism was measured in each condition (right part). The results are presented as the mean of at least three independent experiments ± SD. Statistical analysis was performed using ordinary one-way ANOVA to compare differences between the sicontrol and the other two groups or among all the experimental groups **(E)**. Statistical significance is denoted as follows: *P < 0.05, **P < 0.01, ***P < 0.001, ****P < 0.0001, NS (non-significant).

### Inhibition of USP39 Triggers G2/M Cell Cycle Arrest and Apoptosis in Multiple Myeloma Cells

To elucidate the underlying mechanisms driving the inhibitory effects of USP39 suppression on MM cell growth and colony formation, we conducted a comprehensive analysis of cell cycle progression using flow cytometry over a time course ranging from 72 to 120 hours (**Figure 4A**). Our findings revealed that compared to control siRNA, USP39 siRNA induced a reduction in the G0/G1-phase population (57.20% vs. 35.61% at 72h) and a concurrent increase in the G2/M-phase population (23.78% vs. 39.12% at 72h). Notably, the percentage of cells in the S-phase remained unaffected by USP39 depletion. Additionally, we observed a significant rise in the Sub-G1 phase population (representing dead cells) following USP39 siRNA transfection compared to control siRNA transfection (6.27% vs. 22.05% at 120h). Nocodazole served as a positive control for inducing cell cycle arrest in the G2/M phase. Collectively, these results underscore the role of USP39 depletion in causing G2/M-phase arrest in MM cells. Furthermore, to elucidate the apoptotic process triggered by USP39 depletion, we examined apoptosis in MM cells. Flow cytometry analysis revealed that compared to control siRNA, siUsp39 increased the proportion of Annexin V+/DAPI-(18.37% vs. 5.3%) and Annexin V+/DAPI+ (45.78% vs. 19.89%) populations, indicative of cells in early and late apoptosis, respectively (**Figure 4B**). These findings were corroborated by immunoblotting, which detected cleavage of Parp and Caspase 3 following USP39 depletion (**Figure 4C**). Moreover, the increase in caspase activation induced by USP39 depletion was confirmed by measuring the catalytic activity of caspase 9 initiator and caspase 3 effector (**Figure 4D-E**). Throughout these experiments, the proteasome inhibitor BTZ was employed as a positive inducer of apoptosis. In summary, our findings underscore that USP39 inhibition induces G2/M cell cycle arrest and apoptosis in MM cells.

**Figure 4:**
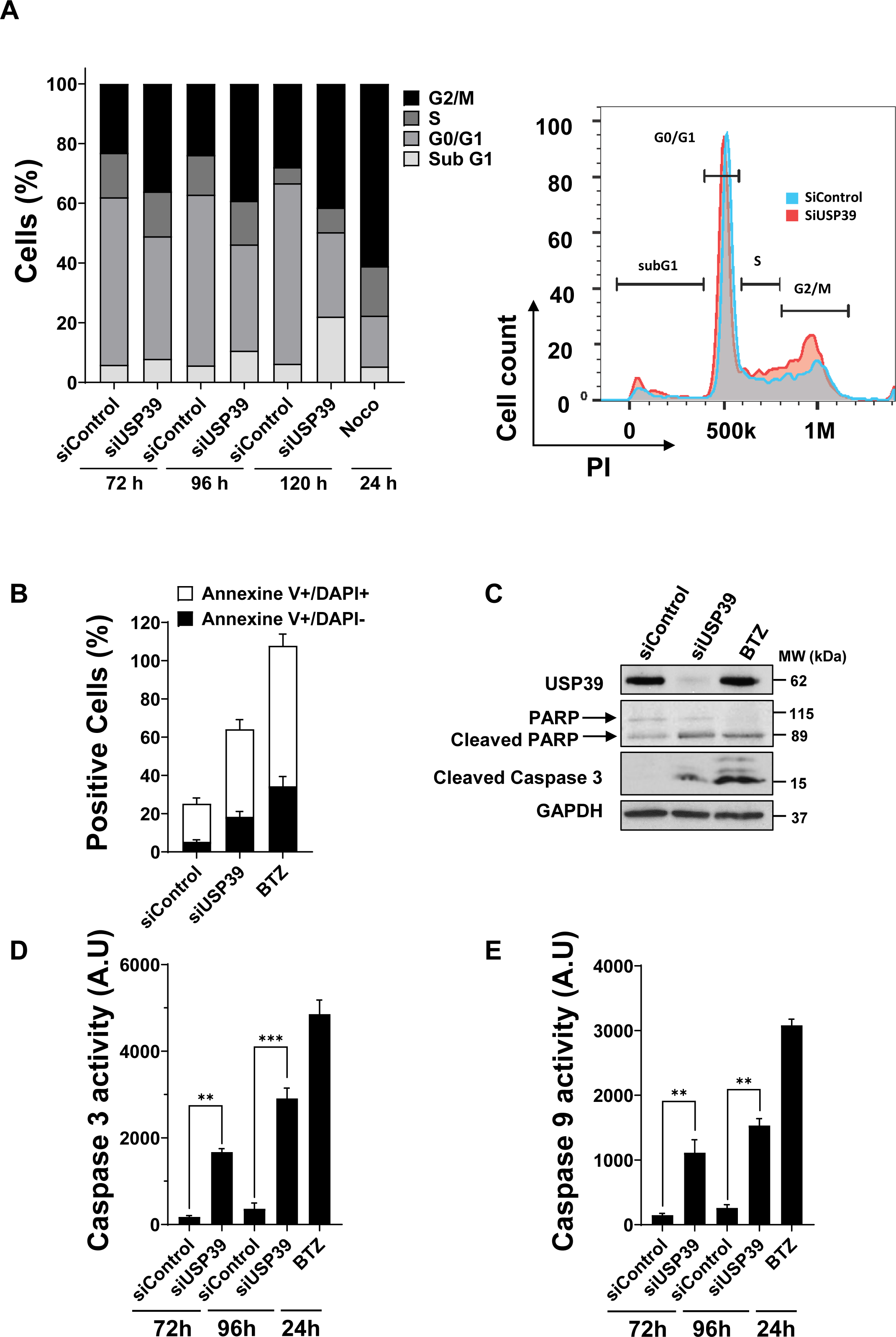
Inhibition of USP39 Triggers G2/M Cell Cycle Arrest and Apoptosis in Multiple Myeloma Cells. **(A)** OPM2 cells were transfected with control or USP39 siRNAs for 72 hours, 96 hours or 120 hours. In parallel, cells were stimulated with nocodazole (1 µg/ml) for 24 hours to block the cells in G2/M phase. Cell cycle distribution was examined by flow cytometry, and percentage of cells in each phase is indicated (right). A representative flow cytometry profile of cells transfected with control (blue area) or USP39 siRNAs (red area) for 96 hours (left). **(B)** OPM2 cells were transfected with either control or single USP39 siRNAs for 96 hours or stimulated with BTZ for 48 hours. Then, cells were stained by Annexin and PI and analyzed by flow cytometry. % of apoptotic cells (annexin V+/DAPI-) and dead cells (annexin V+/DAPI+) are represented in grey and black respectively. **(C)** Lysates from these cells were subjected to immunoblots using USP39, PARP, cleaved caspase 3 and GAPDH antibodies as a loading control. **(D,E)** OPM2 cells were transfected with either control or single USP39 siRNAs for 72 hours and 96 hours, or stimulated with BTZ for 48 hours. Then, cells lysates were subjected to caspase 3 **(D)** and caspase 9 **(E)** assays. The results are presented as the mean of at least three independent experiments ± SD. Statistical analysis was performed using ordinary one-way ANOVA to compare differences between all the experimental groups. Statistical significance is denoted as follows: *P < 0.05, **P < 0.01, ***P < 0.001, ****P < 0.0001, NS (non-significant).

### inhibition of USP39 resensitizes cells to BTZ USP39 Inhibition Reverses Bortezomib Resistance in MM Cells

Bortezomib (BTZ, PS341, Velcade) stands as the pioneering proteasome inhibitor approved for treating MM, eliciting remarkable response rates in both relapsed/refractory and newly diagnosed MM patients[29]. Nonetheless, a subset of MM patients either fails to respond to BTZ therapy or experiences transient responses followed by relapse. [30].

To evaluate the impact of targeting USP39 on MM cell response to BTZ, we conducted experiments using BTZ-sensitive (U266) and -resistant (U266R) MM cell lines, the latter being previously generated in our laboratory (**Figure 5**) [27]. Consistent with prior reports, BTZ exhibited dose-dependent inhibition of cell metabolism, with an IC50 of approximately 5nM in U266 parental cells, while U266R cells showed resistance, with no discernible IC50 at 100nM (**Figure 5A**). Subsequently, both cell lines were transfected with either control siRNA or one of three USP39 siRNAs for 72 hours. Remarkably, USP39 siRNAs reduced cell metabolism by 60% in both cell lines, suggesting that targeting USP39 overcomes proteasome inhibitor resistance (**Figure 5B**). To ascertain whether USP39 depletion could resensitize myeloma cells to BTZ, parental U266 and U266R cells were transfected with the aforementioned USP39 siRNAs for 72 hours and then exposed to escalating doses of BTZ (ranging from 1 to 100nM) for 24 hours (**Figure 5C**). As anticipated, we observed an additive effect of the USP39 siRNA/BTZ combination in parental cells. Intriguingly, the decline in mitochondrial metabolism induced by USP39 depletion was potentiated by BTZ treatment in U266R cells. Collectively, these experiments underscore the capability of targeting USP39 to overcome BTZ resistance in MM cells.

**Figure 5:**
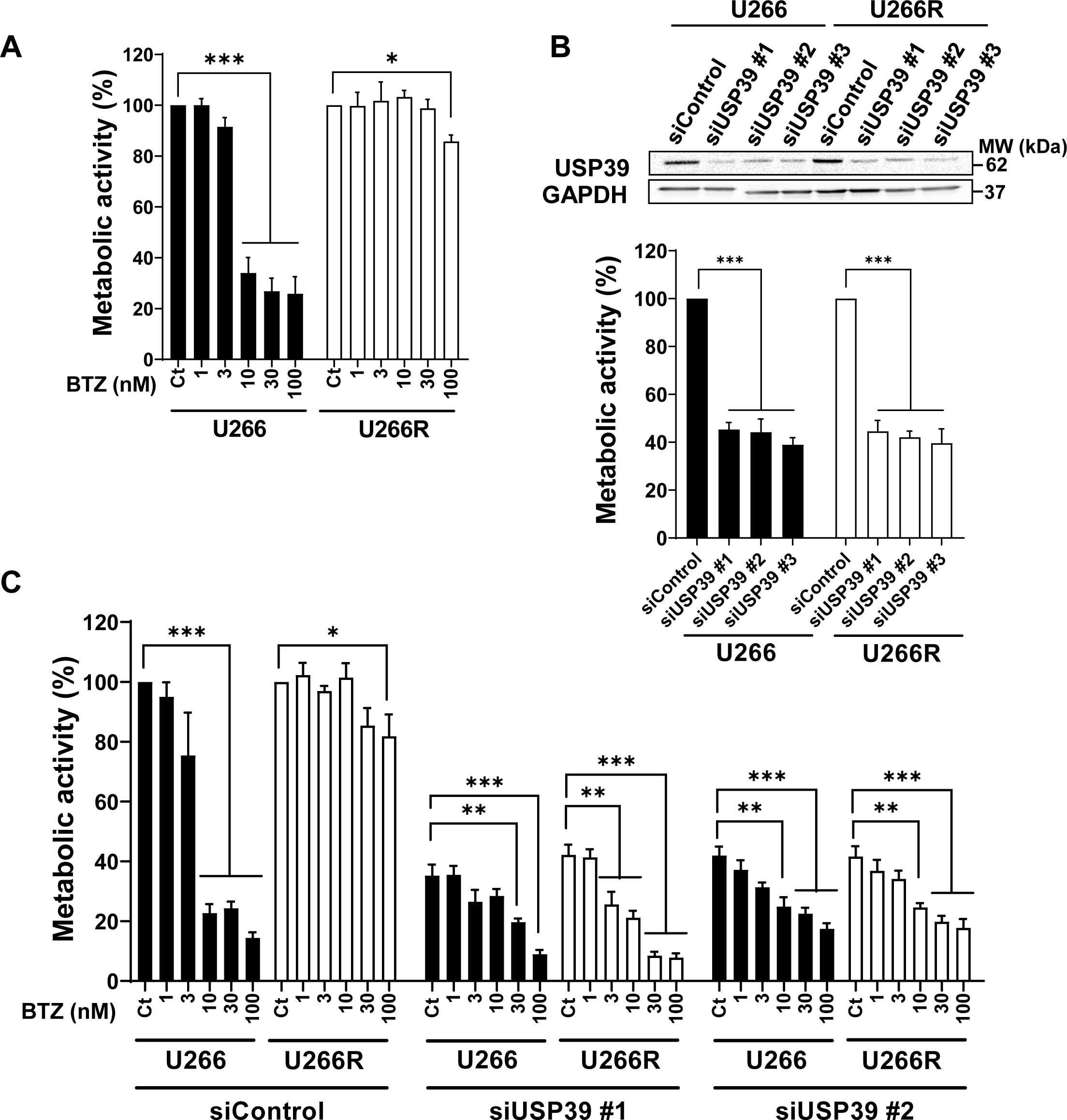
USP39 Inhibition Reverses Bortezomib Resistance in MM Cells. **(A)** U266 cells and its BTZ-resistant counterpart U266R were stimulated with increased concentrations of BTZ (1, 3, 10, 30 and 100 ng/ml) for 24 hours, and cell metabolism was measured by XTT assay. **(B)** U266 and U266R cells were transfected with either control or three different single USP39 siRNAs (#1, #2 and #3) for 72 hours. USP39 silencing was confirmed by immunoblots using USP39 and GAPDH antibodies (top). In parallel, cell metabolism was measured by XTT assay (bottom). **(C)** U266 and U266R cells were transfected with either control or two different single USP39 siRNAs (#1 and #2) for 72 hours. Then both cells were stimulated with increased concentrations of BTZ (1, 3, 10, 30 and 100 ng) for 24 hours and cell metabolism was measured by XTT assay. The results are presented as the mean of at least three independent experiments ± SD. Statistical analysis was performed using ordinary one-way ANOVA to compare differences among all the experimental groups. Statistical significance is denoted as follows: *P < 0.05, **P < 0.01, ***P < 0.001, ****P < 0.0001, NS (non-significant).

### USP39 enhances ZEB1 Protein Stability in a proteasome-dependent manner

To gain deeper insights into the mechanism underlying USP39 targeting, we assessed the expression levels of some of its known substrates. OPM2 cells were transfected with either control or USP39 siRNAs for 48h, 72h, or 96h, and the expression of USP39, ZEB1, SP1, CHK2, and STAT1 was evaluated by immunoblot analysis (**Figure 6A**) and quantified **(Figure S3)**. Among these substrates, only ZEB1 expression exhibited a significant decrease upon USP39 depletion. Conversely, we demonstrated that overexpression of USP39 in MM KMM1 cells led to an increase in ZEB1 expression (**Figure 6B**). Moreover, treatment with the protein translation inhibitor cycloheximide resulted in a reduction in the half-life of the ZEB1 protein, whereas overexpression of USP39 partially restored its stability (**Figure 6C**). To further elucidate the mechanism by which USP39 stabilizes ZEB1 expression, we transfected USP39 siRNA into OPM2 cells and subsequently treated them with the proteasome inhibitor MG132 (**Figure 6D**). Immunoblot analysis revealed that MG132 partially rescued the degradation of ZEB1 following USP39 depletion. To confirm that USP39 regulates ZEB1 at the protein level, we showed by q-PCR that USP39 siRNA does not decrease ZEB1 mRNA and vice versa (**Figure S4**). These findings collectively suggest that USP39 regulates ZEB1 protein stability in a proteasome-dependent manner.

**Figure 6:**
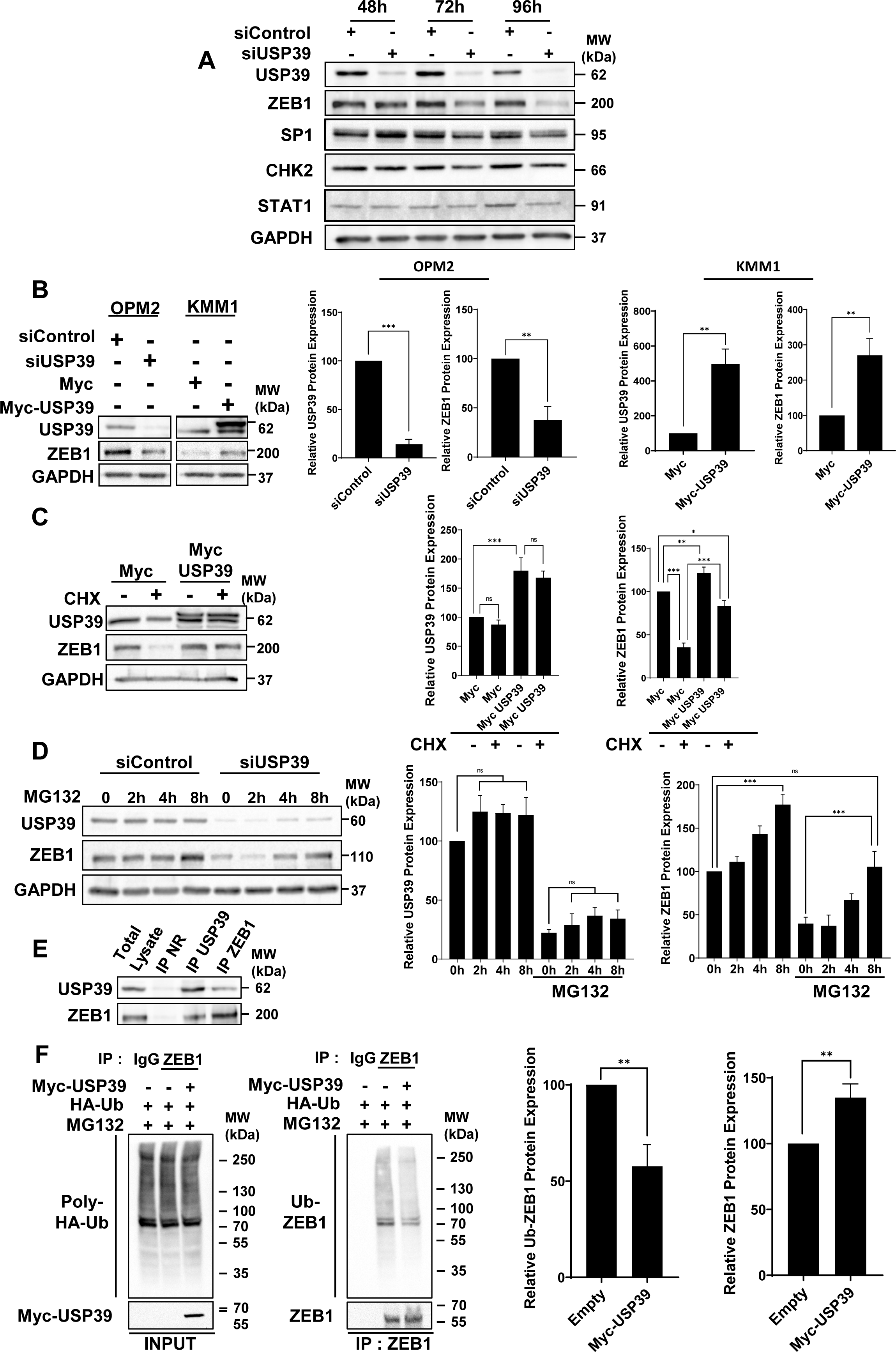
USP39 Deubiquitinates ZEB1 Protein in Multiple Myeloma Cells. **(A)** OPM2 cells were transfected with either control or USP39 siRNAs for 48 hours, 72 hours or 96 hours. Lysates were subjected to immunoblots using USP39, ZEB1, SP1, CHK2, STAT1 and GAPDH antibodies. **(B)** USP39 was transiently silenced or overexpressed in OPM2 and KMM1 cells respectively. Then, immunoblots were performed using USP39, ZEB1 and GAPDH antibodies (left) and protein quantifications were determined (right). **(C)** KMM1 cells were transfected with plasmids encoding either the Myc-tag or the USP39-Myc-tag proteins. After 48 hours, cells were stimulated with cycloheximide (CHX) at 10 µM for 24 hours. Lysates were subjected to immunoblots using USP39, ZEB1, and GAPDH antibodies and protein quantification was determined. **(D)** OPM2 cells were transfected with either control or USP39 siRNAs for 72h. Then cells were stimulated for 2 hours, 4 hours or 8 hours with the proteasome inhibitor MG132 at 1 µM. Lysates were subjected to immunoblots using USP39, ZEB1 and GAPDH antibodies. Protein quantification was determined. **(E)** Lysates from OPM2 cells were subjected to co-immunoprecipitation experiments using either non relevant (NR), USP39 or ZEB1 antibodies. Immunoblots was performed to visualize complexes using USP39 and ZEB1 antibodies. **(F)** KMM1 cells were transfected for 48 hours with HA-ub plasmid in the presence or in the absence of Myc-USP39 plasmid. Then cells were treated with MG132 at 1 µM for 8 hours and lysates were subjected to immunoprecipitation using non relevant IgG or ZEB1 antibodies. Inputs were immunoblotted with HA and Myc antibodies to visualize poly-HA-Ub and USP39 respectively. IPs products were immunoblotted with HA and ZEB1 antibodies to visualize Ub-ZEB1 complex and immunoprecipitated ZEB1. The graph represents Ub-ZEB1 quantification. The results are presented as the mean of at least three independent experiments ± SD. Differences between the two experimental groups were analysed using the Unpaired Student’s t-test **(B, F)**. Statistical analysis was performed using ordinary one-way ANOVA to compare differences among all the experimental groups **(C, D)**. Statistical significance was denoted as follows: *P < 0.05, **P < 0.01, ***P < 0.001, ****P < 0.0001, NS (non-significant).

### USP39 Deubiquitinates ZEB1 Protein in Multiple Myeloma Cells

To elucidate whether USP39 modulates the ubiquitination status of ZEB1, we initially examined the interaction between the USP39 enzyme and its substrate ZEB1. Co-immunoprecipitation assays demonstrated the formation of a complex between endogenous USP39 and ZEB1 proteins in OPM2 cells (**Figure 6E**). To assess the functional relevance of this interaction, deubiquitination assays were conducted. (**Figure 6F**). KMM1 cells were transfected with HA-ubiquitin plasmid in the presence or absence of Myc-USP39 plasmid and treated with MG132 to inhibit proteasomal degradation of HA-ubiquitin proteins. Immunoprecipitation of endogenous ZEB1 followed by immunoblotting with HA and Myc antibodies revealed transfected poly-HA-Ubiquitin and exogenous Myc-USP39, respectively (left part). Immunoblotting of immunoprecipitated products with HA and ZEB1 antibodies demonstrated the presence of HA-Ubiquitin-ZEB1 complex and endogenous ZEB1, respectively (right part). Consistently, protein quantification revealed that exogenous USP39 transfection led to an increase in endogenous ZEB1 expression coupled with a decrease in its ubiquitination level. Collectively, these findings suggest that in MM cells, USP39 regulates the stability of ZEB1 in a proteasome-dependent manner through its deubiquitination activity.

### USP39 Promotes In Vitro Transmigration of MM Cells

It is well-established that the transcription factor ZEB1 plays a pivotal role in promoting tumor invasion and migration by orchestrating the epithelial-to-mesenchymal transition (EMT) [31]. Given USP39’s role in regulating ZEB1 stability, we investigated the impact of USP39 depletion on the migratory capacity of MM cells (**Figure 7A**). Initially, we assessed the expression levels of EMT markers in OPM2 cells depleted for either USP39 or ZEB1 for 48 hours. ZEB1 depletion resulted in reduced expression of β-Catenin, N-Cadherin, and Vimentin proteins, while USP39 depletion led to decreased expression of ZEB1, β-Catenin, and N-Cadherin, consistent with EMT inhibition. E-Cadherin expression was not detected in OPM2 cells. Subsequently, we demonstrated that depletion of both ZEB1 and USP39 attenuated transmigration of OPM2 cells, as evidenced by Boyden chamber assays, compared to non-depleted OPM2 cells (**Figure 7A lower left)**. Moreover, assessment of OPM2 cell viability under the same conditions revealed that neither ZEB1 nor USP39 depletion affected cell viability at 48 hours, suggesting that USP39 depletion reduces MM cell transmigration independently of late cell death (**Figure 7A lower right)**. To ascertain that the observed migratory effect resulting from USP39 depletion is mediated by the regulation of ZEB1, we conducted a complementation approach. Our results showed that the inhibition of migration induced by USP39 depletion was partially restored by exogenous transfection of ZEB1 (**Figure 7B**).

**Figure 7:**
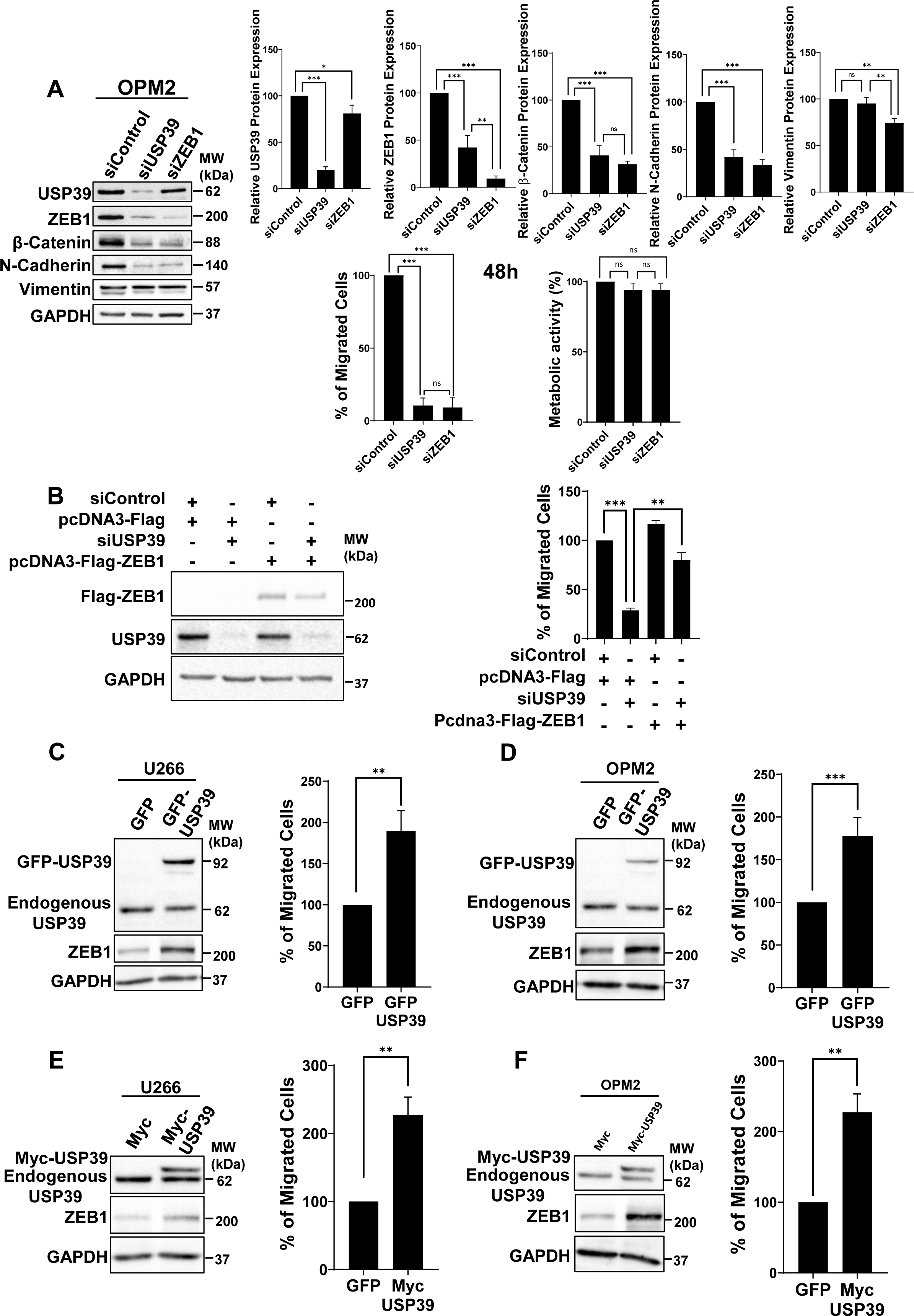
USP39 Promotes In Vitro Transmigration of MM Cells. **(A)** OPM2 cells were transfected with control, USP39 or ZEB1 siRNAs for 48 hours. Then, immunoblots were performed using USP39, ZEB1, β-Catenin, N-Cadherin, Vimentin and GAPDH antibodies (left) and protein quantifications were determined (right). In parallel, the metabolic activity (lower left) and the migration of the cells (lower right) were measured under the same conditions. **(B)** KMM1 cells were either transfected with Control or USP39 siRNAs for 24 hours. Then cells were transfected with pcDNA3-Flag or pcDNA3-Flag-USP39 vectors. After 72 hours, lysates from these cells were subjected to immunoblots using USP39, Flag-Tag or GAPDH, antibodies (left part). After 96 hours of transfections, the migration capacity of cells was measured by Boyden chamber assays (right part). **(C and D)** U266 **(C)** and OPM2 **(D)** cells were stably transduced with lentiviral particles encoding GFP or GFP-USP39 and were subjected to immunoblots using USP39, ZEB1 or GAPDH antibodies. **(E, F)** U266 **(E)** and OPM2 **(F)** cells were stably transfected with lentiviral particles encoding Myc or Myc-USP39 and were subjected to immunoblots using USP39, ZEB1 or GAPDH antibodies. In parallel, the migration capacity of corresponding cells was measured by Boyden chamber assays. The results are presented as the mean of at least three independent experiments ± SD. Statistical analysis was performed using ordinary one-way ANOVA to compare differences among all the experimental groups **(A-B)**. Differences between the GFP and GFP-USP39 or Myc-USP39 were analysed using the Unpaired Student’s t-test **(C-F)**. Statistical significance was denoted as follows: *P < 0.05, **P < 0.01, ***P < 0.001, ****P < 0.0001, NS (non-significant).

Furthermore, to reinforce our findings, U266 and OPM2 cells were stably transfected with lentiviral particles encoding either GFP or GFP-USP39 (**Figure 7C-D**) or Myc and Myc-USP39 (**Figure 7E-F**). The nuclear localization of the GFP-USP39 form was confirmed by immunofluorescence. **(Figure S5)**. Immunoblots revealed overexpression of both forms of USP39 in OPM2 and U266 cells, accompanied by increased expression of ZEB1. Finally, Boyden chamber assays demonstrated that OPM2 and U266 cells overexpressing either the GFP-USP39 or Myc-USP39 form exhibited enhanced migratory capacity. Collectively, our gain-of-function and loss-of-function experiments strongly suggest that targeting USP39 modulates the migratory capacity of myeloma cells through the regulation of ZEB1 expression.

### USP39 Enhances Metastasis in Zebrafish: Implications for MM Progression

Cell metastasis, a is complex process involving several steps, including cell invasion, egress, entry into the circulation and specific homing to predetermined distant tissues. In hematological cancers, including MM, bone marrow (BM) dissemination significantly impacts disease aggressiveness and patient outcomes. Zebrafish larvae provide the opportunity to observe the behavior of transplanted tumor cells using high-resolution in vivo imaging techniques and to rapidly analyze the metastatic behavior of human MM cells [32]. As we have shown that USP39 promotes the EMT process and MM cell transmigration *“in vitro”*, Zebrafish xenograft model was used to determine whether USP39 overexpression could promote cell migration and invasion “*in vivo*” (**Figure 8**). Injection of OPM2 cells stably overexpressing USP39 (Myc-USP39) or control cells (Myc) into the perivitelline space of zebrafish larvae allowed for real time visualization of local and distant metastatic events using fluorescence microscopy after 2 days. Our findings reveal a striking difference in the metastatic behavior of USP39-overexpressing cells compared to controls. While both cell types initially remained equivalently confined to the perivitelline space at day 0 post-injection (**Figure 8A-B**), by day 2, a clear divergence was observed. USP39-overexpressing cells exhibited enhanced migration and invasion, resulting in the formation of metastases both locally and at a distance from the injection site. This was evidenced by the presence of fluorescent foci in the body and tail regions of the larvae, indicating successful colonization of distant tissues by USP39-overexpressing MM cells (**Figure 8A left)**. Importantly, an enlarged photo of the larvae’s tail highlighted the enhanced metastatic potential of USP39-overexpressing cells, underscoring the pro-metastatic role of USP39 in MM progression (**Figure 8A right)**. A quantitative analysis further corroborated these observations, revealing a significant increase in the number of distant metastatic foci in larvae injected with USP39-overexpressing cells (**Figure 8C**). Fluorescence quantification indicated that the area of expansion of local metastases was also significantly increased in cells overexpressing USP39 (**Figure 8D**).

**Figure 8:**
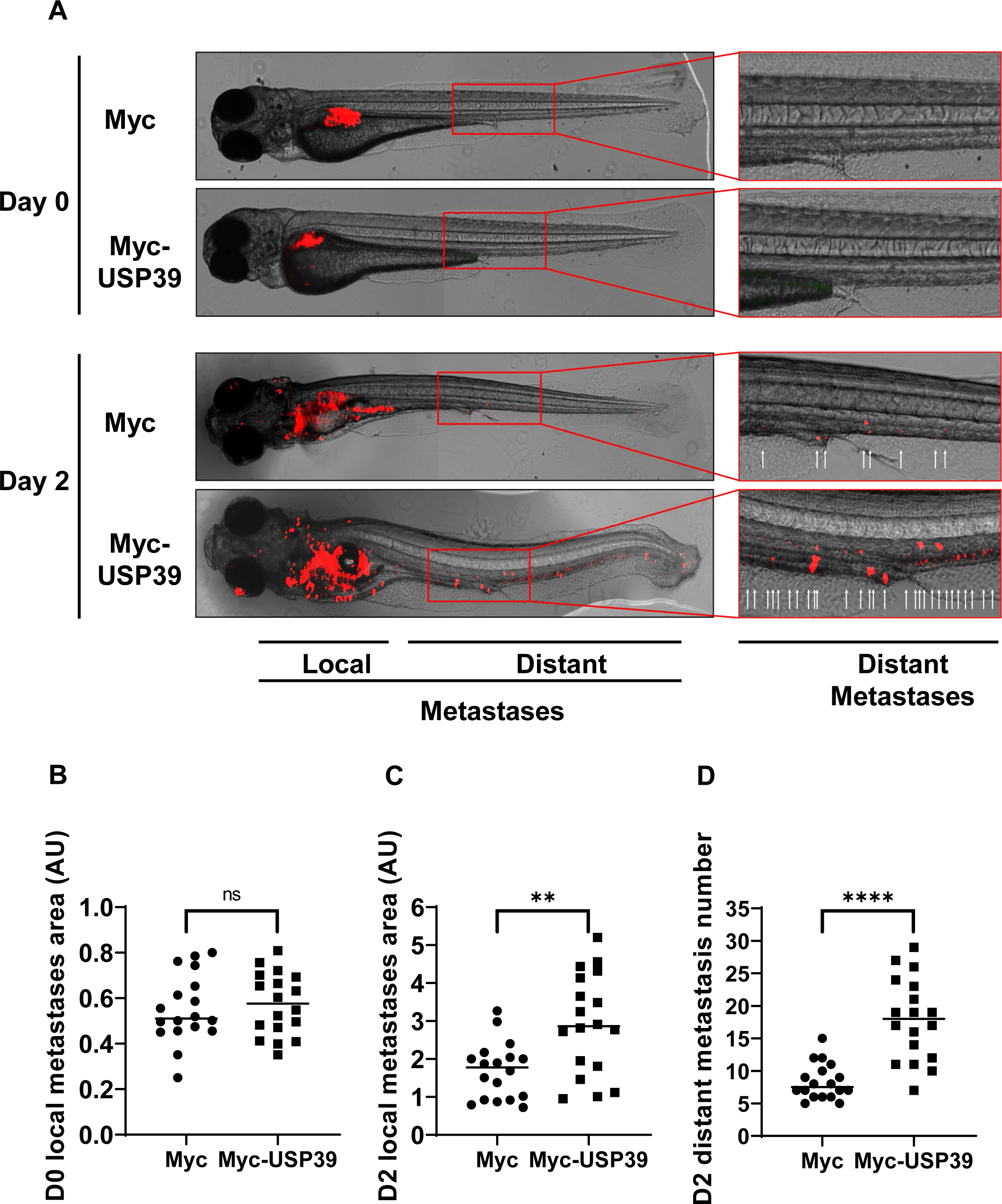
USP39 Enhances Metastasis in Zebrafish: Implications for MM Progression. **(A-D)** Zebrafish embryos (N = 36) were injected with U266 cells stably infected with lentiviral particles encoding either Myc-tag or Myc-USP39 proteins (labeled with red DiD) into the perivitelline space. Zebrafish embryos were monitored at Day 0 and Day 2 for tumor metastases using a fluorescent microscope. **(A) (Left)** Representative images of local and distant metastases are shown. **(Right)** Magnification of images representing distant metastases. Arrows indicate metastases. Quantification of the area of local metastases at Day 0 **(B)** and Day 2 **(C)** of Myc-tag and Myc-USP39 embryos. **(D)** Quantification of the area of distant metastases at Day 2 of Myc-tag and Myc-USP39 embryos. The Mann-Whitney U test was performed to compare the average measurement between Myc-tag and Myc-USP39 embryos. (U = 151, *P < 0.05, **P < 0.01, ***P < 0.001, ****P < 0.0001, NS (non-significant).

These findings provide valuable insights into the mechanisms underlying MM metastasis and underscore the importance of the USP39/ZEB1 axis as a potential therapeutic target. By elucidating the role of USP39 in promoting MM metastasis in vivo, our study opens avenues for further research aimed at developing targeted interventions to inhibit metastatic spread and improve patient outcomes in MM.

Collectively, we have identified an important role of the deubiquitinase USP39 suggesting that the USP39/ZEB1 axis might be pursued as a potential diagnostic and therapeutic target in MM cancer.

## DISCUSSION

The dysregulation of DUBs has emerged as a significant mechanism in the pathogenesis of hematologic malignancies [12], with different roles observed in different contexts, including oncogenic and tumor suppressor functions. Several potent inhibitors targeting oncogenic DUBs have shown promise in preclinical models, and clinical trials [33]. In the realm of MM, various DUBs including USP7, USP9X, USP14, PSMD14 and USP10 have been identified as potential therapeutic targets in MM, yet the comprehensive landscape of DUB involvement in hematologic malignancies, including MM, remains largely unexplored.

Our study unveils the critical role of USP39 as a survival factor for MM cells, using a comprehensive loss-of-function approach to elucidate its function. Among the tested DUBS, inhibition of USP39 leads to the most significant cytotoxic effects in MM cells, suggesting a dependency of MM cells on USP39 for their survival. This observation was further validated across three different MM cell lines, reinforcing the essential role of USP39 in MM pathogenesis. Mechanistically, our study delineates the molecular pathways through which USP39 exerts its pro-survival effects in MM cells. We found that USP39 depletion by siRNA induces a cell cycle arrest at the a G2/M phase, indicative of its crucial role in cell proliferation. These results are consistent with Jian Yuan’s team, which demonstrated that USP39 regulates the DNA damage response and chemo-radiation resistance by deubiquitinating and stabilizing CHK2 (checkpoint kinase 2) [22]. Although we were unable to detect a significant modulation of CHK2 kinase upon USP39 depletion in the OPM2 line (**Figure 6A**), the study of this regulatory axis merits further investigation.

Taken together, our results based on loss-of-function experiments and those reported in the literature confirm that among all DUBs, USP39 belongs to the general category of essential genes [34].

To assess the relevance of targeting USP39 as a therapeutic strategy, we evaluated its expression in MM patients. Remarkably, Gene Set Enrichment Analysis (GSEA) unveiled a significant upregulation of USP39 mRNA levels from monoclonal gammopathy of undetermined significance (MGUS) to the MM stage, compared to healthy individuals. The finding imply that the heightened USP39 expression may serve as an early event in the transformation of malignant plasma cells. Notably, the simpler genomic landscape in MGUS, contrasted with the complexity of MM, underscores the significance of USP39 dysregulation in disease progression. Considering the frequent presence of activating mutations in the KRAS proto-oncogene among MM and MGUS patients, with implications for disease prognosis [35], it is noteworthy that USP39 has been implicated in the development of KRAS-driven lung and colon cancers [36]. This suggests a potential role for the USP39/KRAS axis in driving MM progression, thus rendering it a compelling therapeutic target. Immunohistochemistry experiments further confirmed these findings, revealing overexpression of USP39 in the majority of MM patients compared to healthy donors. The inability to detect USP39 expression in healthy individuals suggests that IHC may lack the sensitivity required to detect low levels of USP39.

Proteasome inhibitors and immunomodulatory drugs, with stem cell auto transplantation (ASCT), have significantly improved the treatment landscape for newly diagnosed MM patients and achieved complete remissions in many cases. However, despite these advancements, relapse remains a persistent challenge for a substantial number of patients. Prolonged salvage treatment can often lead to drug resistance, ultimately resulting in fatal outcomes [2]. In this context, genetic and pharmacological inhibition of DUBs such as USP7, RPN11, and USP12 can circumvent the resistance of MM cells to BTZ [16], [37], [38], [39]. To evaluate the feasibility of targeting USP39 as a strategy for treating MM relapse, we used a model of BTZ-resistant MM cells previously established in our laboratory [27]. Depletion of USP39 not only overcame resistance to BTZ but also enhanced the cytotoxic effect of BTZ in resistant cells. Notably, we previously demonstrated that BTZ-resistant cells retained a functional proteasome [27], indicating that depletion of USP39 still facilitated the degradation of its substrates, thereby sensitizing the cells to BTZ.

Despite the absence of a pharmacological inhibitor targeting USP39, our findings indicate that inhibiting USP39 holds promise as a clinically relevant therapeutic strategy, especially for MM patients who exhibit resistance to BTZ therapy.

MM is characterized by the continual spread of MM cells within and outside the BM, with MM progression driven by intricate interactions between the BM microenvironment and MM cells. These interactions influence MM cell migration and dissemination via the bloodstream and create new BM niches [40] highlighting the critical role of the BM microenvironment in MM progression and dissemination [41]. Epithelial-to-mesenchymal transition (EMT) is a fundamental process in embryonic development. It involves changes such as loss of cell-cell adhesion and acquisition of migratory and invasive properties [42]. While extensively studied in solid tumors, the relevance of EMT features in hematological malignancies, including MM has emerged as an area of interest [43]. EMT-like signatures with upregulation of canonical mesenchymal markers has been correlated with poor prognosis in patients suffering from various hematological malignancies including MM (lymphoma, lymphoid and myeloid leukemia) [44]. However, the precise biological implications of EMT markers in hematopoietic cancers remains largely unexplored. For instance, the role of the EMT transcription factor ZEB1, which promotes methylation and downregulation of B-cell lymphoma protein 6 (BCL6), a key transcription associated with a benign profile [45], has not been elucidated until now. In acute myeloid leukemia (AML), poor patient prognosis correlated with the expression of EMT markers, and experimental downregulation of ZEB1 in AML cells inhibited the invasive capacity of this aggressive cancer [46]. Few of these studies have investigated the relationship between the expression of EMT markers and increased cell migration in MM [47], [48], [49]. Surprisingly, no study to date has demonstrated a role for the EMT transcription factor ZEB1 in the migratory potential of MM cells. We are the first to demonstrate that the transcription factor ZEB1 may play a role in the migratory capacity of MM cells and that its expression level is balanced by the deubiquitinase USP39. Our findings shed light on the molecular mechanisms underlying MM cell migration and dissemination, offering new insights into the pathobiology of MM progression. Our results are in perfect agreement with those of Dr. Gang Song’s team, who elegantly demonstrated that USP39 and the E3 ligase TRIM26 balance the level of ZEB1 ubiquitination and thereby determine hepatocellular carcinoma cell proliferation and migration [26]. Although the function of the ubiquitinated TRIM26 protein has not yet been studied in the context of hematological malignancies, the importance of the USP39/ZEB1/TRIM26 axis deserves to be investigated in the context of MM.

Over the past 3 years, the deubiquitination ZEB1 has been intensively studied in various solid tumors. Several DUBs, including USP10, USP18, USP22, USP43, and USP51 have been identified as key regulators of ZEB1 stability, thereby promoting proliferation, migration and invasion of cancer cells across different malignancies [26], [50], [51], [52], [53], [54]. Our study confirms this findings, demonstrating that the regulation of ZEB1 by DUBs constitutes a general mechanism driving cancer cells aggressiveness, which extends to hematological malignancies capable of acquiring EMT-like marker features, such as MM. However, further research is required in MM to determine the most effective therapeutic approach for targeting ZEB1 stability through DUBs.

In conclusion, our study conducted through an RNAi-based synthetic lethal screen, has identified USP39 as a crucial survival factor for MM cells. Elevated expression of USP39 mRNA correlated with shorter survival of MM patients, and strong expression of the USP39 protein is evident in MM plasmocytes compared to healthy individuals. Inhibition of Usp39 reduces clonogenic abilities, induces apoptosis, causes cell cycle arrest, and overcomes BTZ resistance. Gain-of-function experiments further illustrate that USP39 enhances the trans-migratory capacity of MM cells by stabilizing the transcription factor ZEB1.

Overall, our findings underscore the significant role of USP39 in MM cancer progression. Therefore, targeting the USP39/ZEB1 axis holds promise as a novel therapeutic approach in MM, presenting a potential avenue for the development of innovative treatment strategies to improve outcomes for MM patients.

## ABBREVATION

ASCT: stem cell auto transplantation
BM: bone marrow
BTZ: bortezomib
DUB: deubiquitinase
EMT: epithelial-to-mesenchymal transition
GFP: green fluorescent protein
GSEA: Gene Set enrichment analysis
HCC: hepatocellular carcinoma
IMiDs: immunomodulatory drugs
JAAMs: JAB1/PAB1/MPN-domain containing metallo-enzyme
MGUS: monoclonal gammopathy of undetermined significance
MJDs: Machado-Joseph disease protein domain proteases
MM: multiple myeloma
Monoclonal antibodies: mAbs
NR: non relevant
OTUs: ovarian tumor-related proteases
PIs: proteasome inhibitors
siRNA: small interfering RNA
Ub: ubiquitin
Ubis: Ubiquitinases
UCHs: ubiquitin carboxy-terminal hydrolases
UPS: ubiquitin proteasome system
USP39: ubiquitin-specific peptidase 39
USPs: ubiquitin-specific proteases
WT: wild type
ZEB1: Zinc-finger E-box-binding homeobox 1

## ACKNOWLEDGMENT

We kindly thanks Valerie Vouret, Katia Zahaf et Salomé Lalvée for their help for IHC experiments.

## DATA AVAILABILITY STATEMENT

The data that support the findings of this study are available from the corresponding author upon reasonable request.

## SUPPLEMENTAL MATERIAL AND METHODS

### IHC

Immunohistochemistry for CD138 (#760-4248, Roche diagnostics, F. Hoffmann-La Roche, Basel, Switzerland) were performed using Myeloma tissue array (BM291d and BM483b) purchased from TissueArray.com. Antigen retrieval was performed by boiling sections for 10 min in citrate buffer (pH 6.0) and cooling at RT°, followed by blocking of endogenous peroxidase activity with 0.3% H_2_O_2_ for 30 min. The sections were blocked with 2.5% horse serum in TBS solution for 30 min in a humid chamber prior to incubation with anti-USP39 antibody (1/200). Positive cells were detected using an ImmPRESS HRP anti-rabbit detection kit. The immune complexes were visualized using a Peroxidase Substrate DAB kit (Vector) according to the manufacturer’s protocol, and slides were counterstained with hematoxylin.

### qPCR

Total RNA was isolated by using the RNeasy Plus Mini kit (QIAGEN, Hilden, Germany) and quantified by using a Nanodrop 2000 UV visible spectrophotometer. One microgram of mRNA was reverse-transcribed into cDNA by using an QuantiTect Reverse Transcription Kit (QIAGEN), according to the manufacturer’s instructions. Real-time qPCR was performed on cDNA by using StepOnePlus™ Real-Time PCR System (Applied bioscience) and Takyon™ ROX SYBR 2X MasterMix dTTP blue (Eurogentec). Predesigned primer were purchased from Eurogentec (USP39 forward: TTG-GAA-GAG-GCG-AGA-TAA; USP39 reverse: AGG-AGC-ATC-AAT-CAT-CAT-C) and (ZEB1 forward: CAG-CTT-GAT-ACC-TGT-GAA-TGG-G; ZEB1 R: TAT-CTG-TGG-TCG-TGT-GGG-ACT). Fold changes in expression were calculated by the delta Ct method using RPLP0 primers (36B4 forward 5ʹ GGC-CAG-GAC-TCG-TTT-GTA-CC; 36B4 R 5ʹ CAGATTGGCTACCCAACTGTT) gene as an endogenous control for mRNA expression. All fold changes were expressed as normalized to the untreated control. Measurements were done in triplicate.

## SUPPLEMENTAL FIGURE LEGENDS

**Figure S1:**
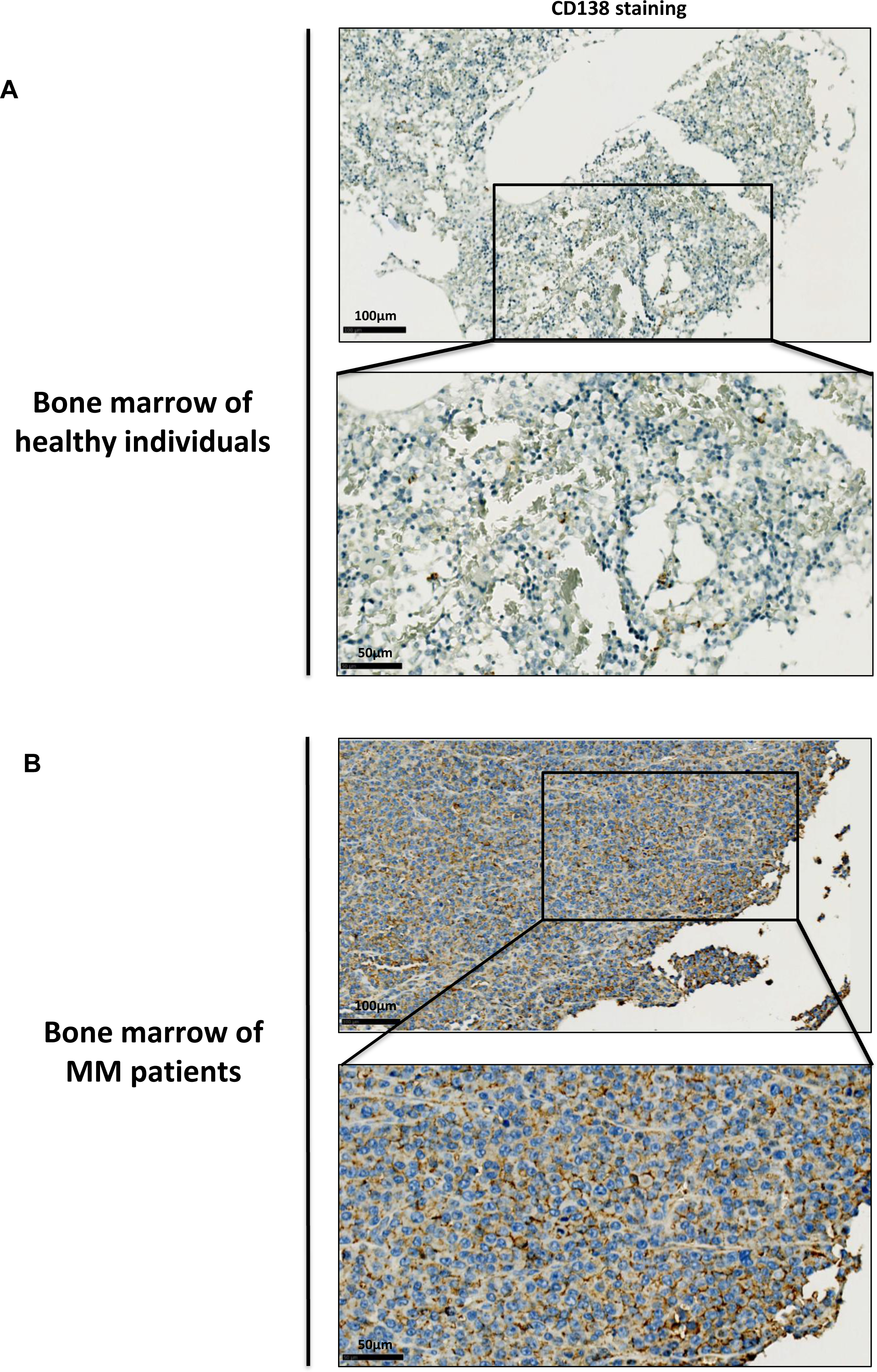
CD138 expression in bone marrow of MM patients. **(A, B)** Representative pictures of CD138 staining from BM tissue sections of healthy donors **(A)** and MM patients **(B)**.

**Figure S2:**
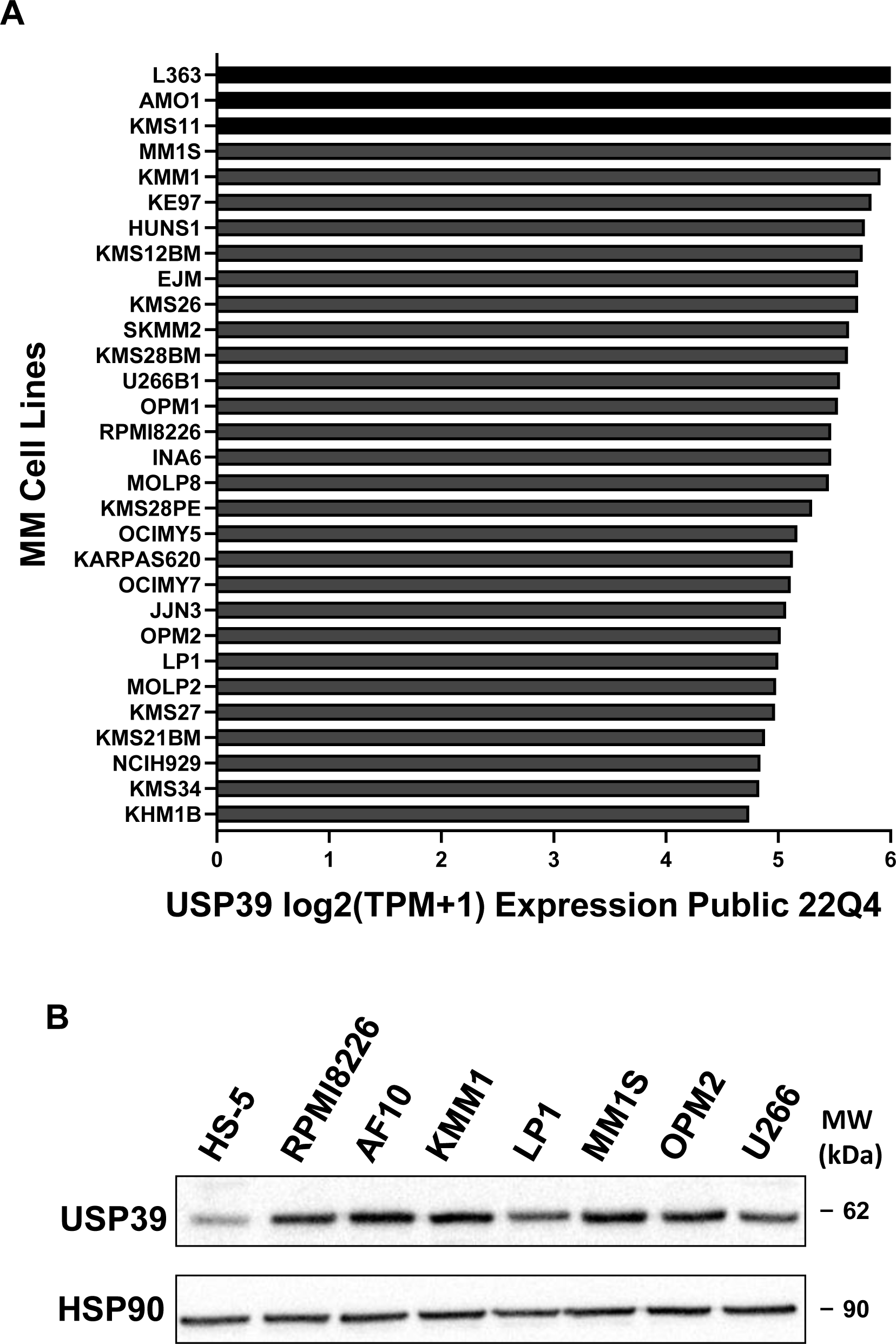
USP39 is widely expressed in MM cell lines. **(A)** Graph represents the mRNA expression levels of different MM cell lines from Depmap portal website. **B)** Cell lysates from different MM cell lines were subjected to immunoblot using USP39 and HSP90 antibodies.

**Figure S3:**
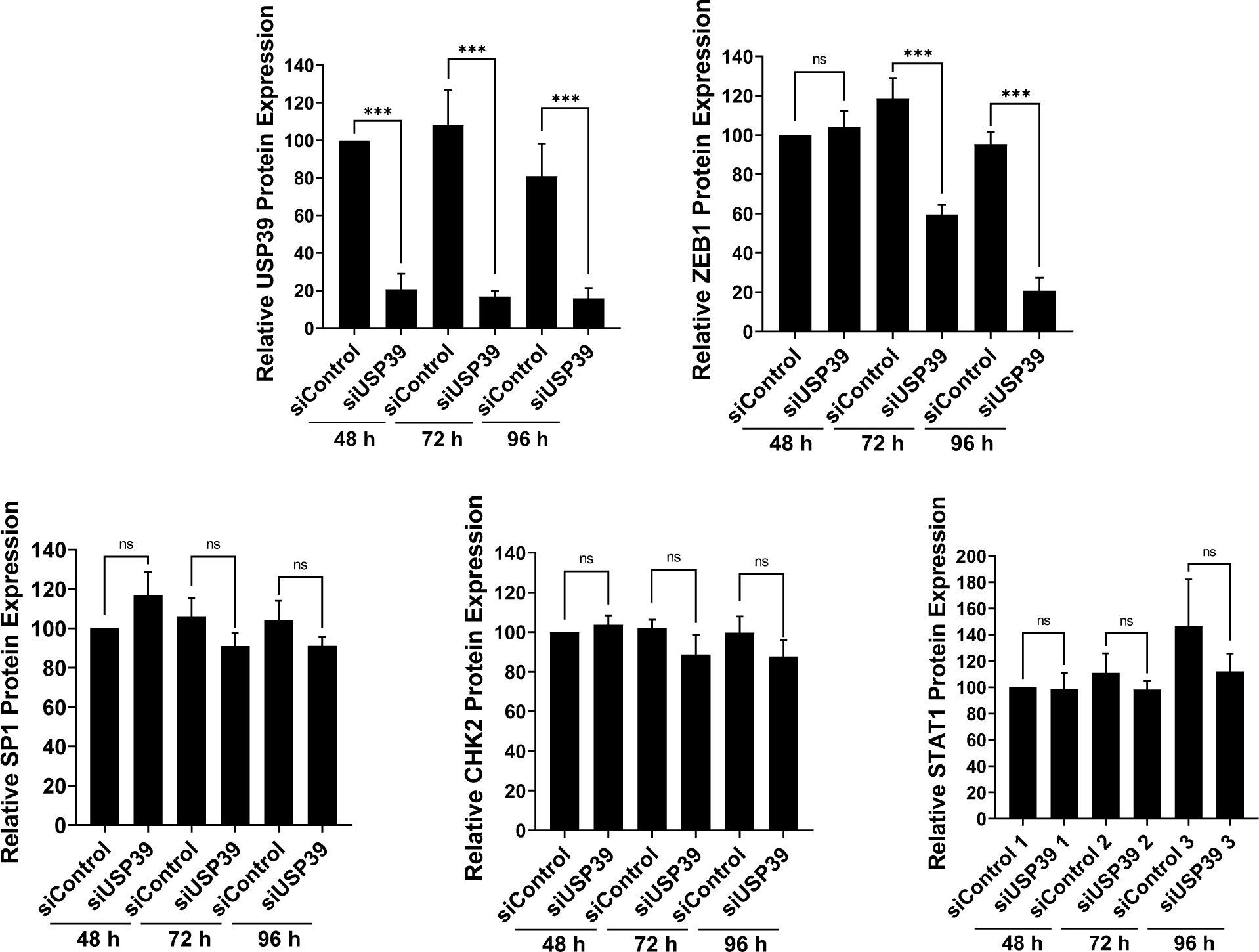
USP39 depletion decreases the expression of ZEB1 over the time. The graphs illustrate the quantification of proteins detected by immunoblot, as shown in Figure 5A. The results are presented as the mean of at least three independent experiments ± SD. Statistical analysis was performed using ordinary one-way ANOVA to compare differences among all the experimental groups. The results are presented as the mean of at least three independent experiments ± SD. Statistical analysis was performed using ordinary one-way ANOVA to compare differences among all the experimental groups. Statistical significance is denoted as follows: *P < 0.05, **P < 0.01, ***P < 0.001, ****P < 0.0001, NS (non-significant).

**Figure S4:**
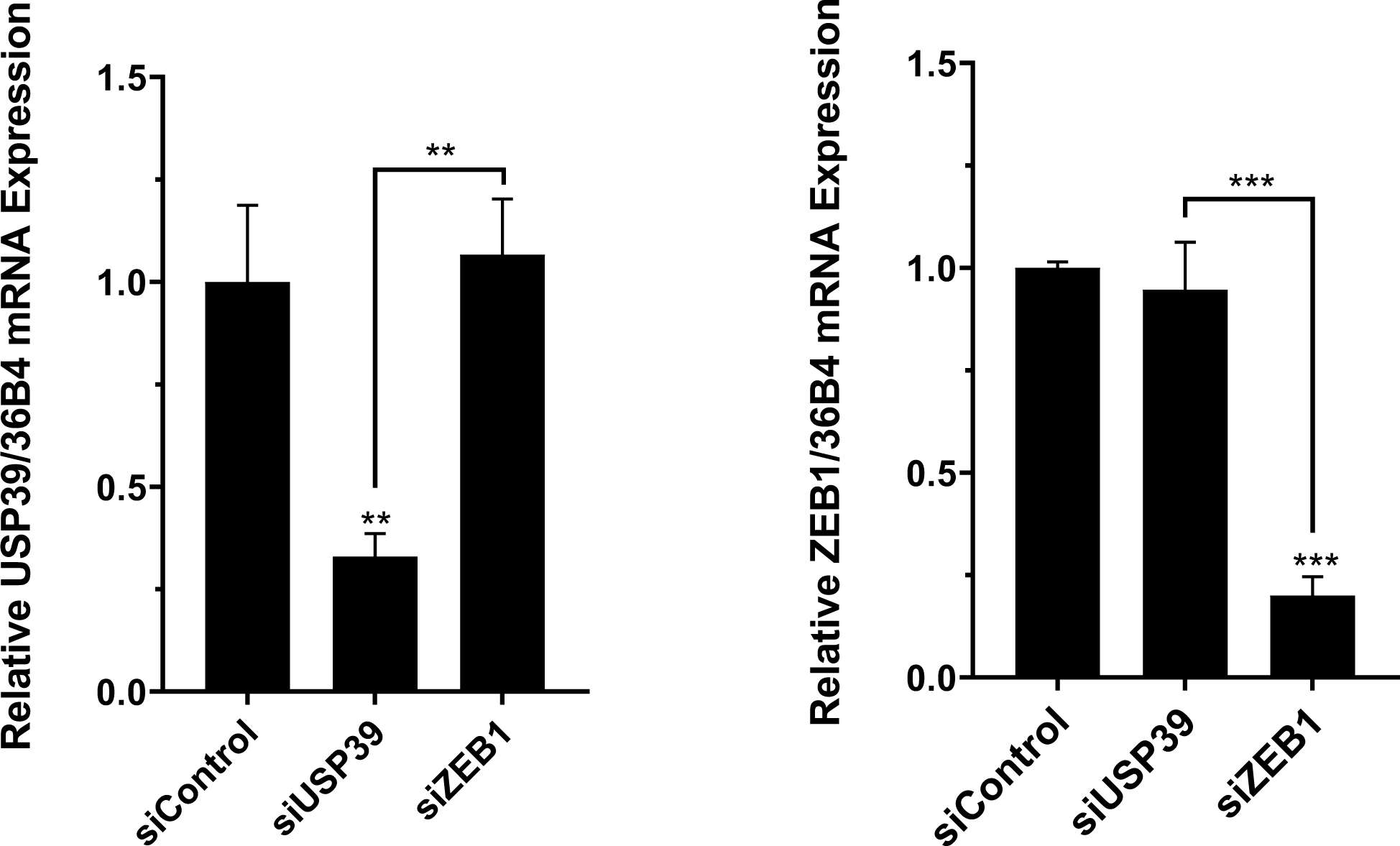
Depletion of USP39 does not decrease ZEB1 mRNA level. The results are presented as the mean of at least three independent experiments ± SD. Statistical analysis was performed using ordinary one-way ANOVA to compare differences among all the experimental groups. Statistical significance is denoted as follows: **P < 0.01, ***P < 0.001.

**Figure S5:**
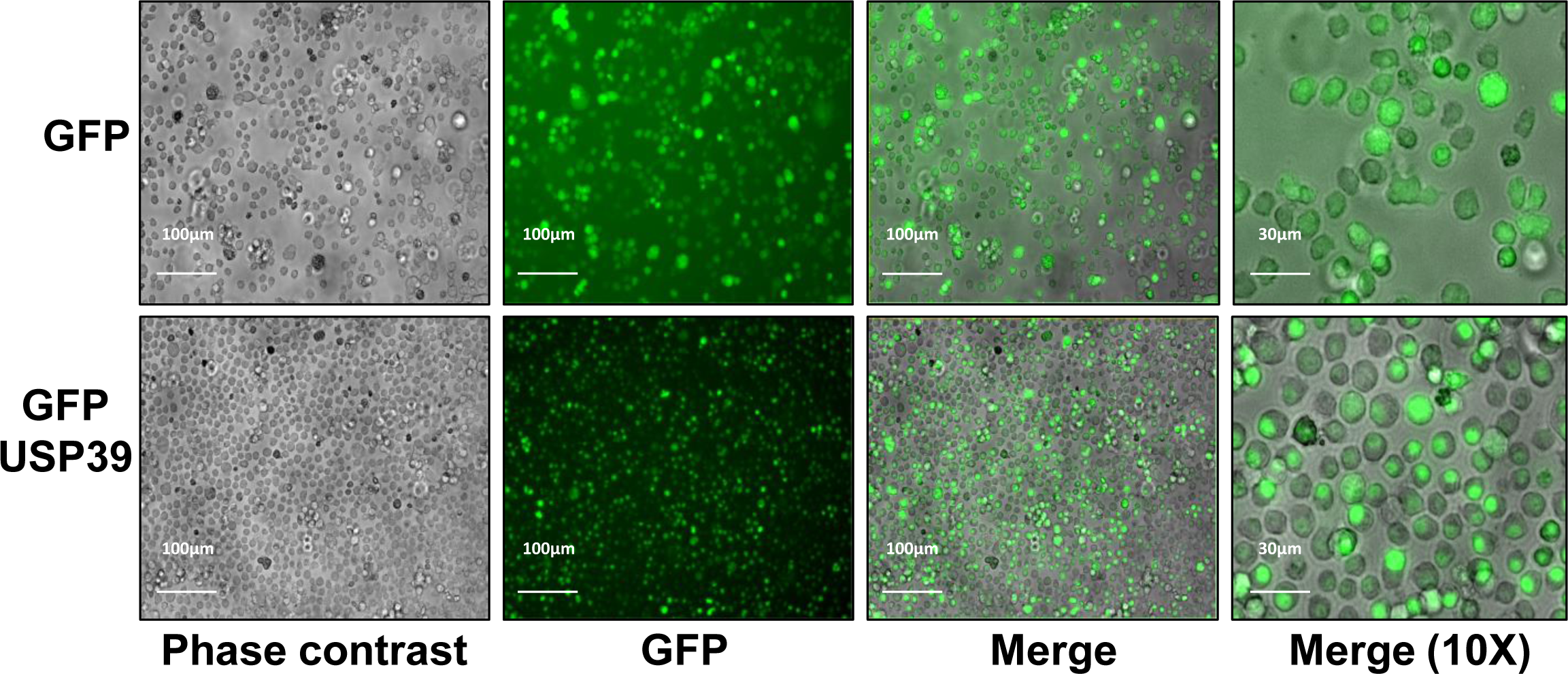
Nuclear localization of USP39 in OPM2 cells. Fluorescence microscopy images of OPM2 cells stably infected with lentiviral particles encoding GFP (top) or GFP-USP39 (bottom) fusion proteins. The right panels represent a 10x magnification.

## REFERENCES

[1] N. W. C. J. van de Donk, C. Pawlyn, et K. L. Yong, « Multiple myeloma », The Lancet, vol. 397, n° 10272, p. 410–427, janv. 2021, doi: 10.1016/S0140-6736(21)00135-5.

[2] V. Pinto, R. Bergantim, H. R. Caires, H. Seca, J. E. Guimarães, et M. H. Vasconcelos, « Multiple Myeloma: Available Therapies and Causes of Drug Resistance », Cancers, vol. 12, n° 2, p. 407, févr. 2020, doi: 10.3390/cancers12020407.

[3] A. H. Bazarbachi, R. Al Hamed, F. Malard, J.-L. Harousseau, et M. Mohty, « Relapsed refractory multiple myeloma: a comprehensive overview », Leukemia, vol. 33, n°10, p. 2343–2357, oct. 2019, doi: 10.1038/s41375-019-0561-2.

[4] D. Nandi, P. Tahiliani, A. Kumar, et D. Chandu, « The ubiquitin-proteasome system », J Biosci, p. 19, 2006.

[5] A. P. K. Wodrich, A. W. Scott, A. K. Shukla, B. T. Harris, et E. Giniger, « The Unfolded Protein Responses in Health, Aging, and Neurodegeneration: Recent Advances and Future Considerations », Front. Mol. Neurosci., vol. 15, p. 831116, févr. 2022, doi: 10.3389/fnmol.2022.831116.

[6] A. A. Sahasrabuddhe et K. S. J. Elenitoba-Johnson, « Role of the ubiquitin proteasome system in hematologic malignancies », Immunol. Rev., vol. 263, n° 1, p. 224–239, janv. 2015, doi: 10.1111/imr.12236.

[7] J. L. Harousseau et M. Attal, « How I treat first relapse of myeloma », Blood, vol. 130, n°8, p. 963–973, août 2017, doi: 10.1182/blood-2017-03-726703.

[8] S. Ito, « Proteasome Inhibitors for the Treatment of Multiple Myeloma », Cancers, vol. 12, n°2, p. 265, janv. 2020, doi: 10.3390/cancers12020265.

[9] K. N. Swatek et D. Komander, « Ubiquitin modifications », Cell Res., vol. 26, n° 4, p. 399–422, avr. 2016, doi: 10.1038/cr.2016.39.

[10] A. Y. Amerik et M. Hochstrasser, « Mechanism and function of deubiquitinating enzymes », Biochim. Biophys. Acta, p. 19.

[11] T. E. T. Mevissen et D. Komander, « Mechanisms of Deubiquitinase Specificity and Regulation », Annu. Rev. Biochem., vol. 86, n° 1, p. 159–192, juin 2017, doi: 10.1146/annurev-biochem-061516-044916.

[12] H. Lei, J. Wang, J. Hu, Q. Zhu, et Y. Wu, « Deubiquitinases in hematological malignancies », Biomark. Res., vol. 9, n° 1, p. 66, déc. 2021, doi: 10.1186/s40364-021-00320-w.

[13] Y. Xu et al., « Targeting the Otub1/c-Maf axis for the treatment of multiple myeloma », Blood, vol. 137, n° 11, p. 1478–1490, mars 2021, doi: 10.1182/blood.2020005199.

[14] S. Hussain, T. Bedekovics, M. Chesi, et P. L. Bergsagel, « UCHL1 is a biomarker of aggressive multiple myeloma required for disease progression », p. 15.

[15] M. Schwickart, « Deubiquitinase USP9X stabilizes MCL1 and promotes tumour cell survival », vol. 463, p. 6, 2010.

[16] R. Franqui-Machin et al., « Destabilizing NEK2 overcomes resistance to proteasome inhibition in multiple myeloma », J. Clin. Invest., vol. 128, n° 7, p. 2877–2893, juill. 2018, doi: 10.1172/JCI98765.

[17] X. Dong et al., « USP39 promotes tumorigenesis by stabilizing and deubiquitinating SP1 protein in hepatocellular carcinoma », Cell. Signal., vol. 85, p. 110068, sept. 2021, doi: 10.1016/j.cellsig.2021.110068.

[18] « Wang et al. - 2013 - Lentivirus-mediated inhibition of USP39 suppresses.pdf ».

[19] Z. Xing, F. Sun, W. He, Z. Wang, X. Song, et F. Zhang, « Downregulation of ubiquitin-specific peptidase 39 suppresses the proliferation and induces the apoptosis of human colorectal cancer cells », Oncol. Lett., févr. 2018, doi: 10.3892/ol.2018.8061.

[20] J. Yuan et al., « Knocking down USP39 Inhibits the Growth and Metastasis of Non-Small-Cell Lung Cancer Cells through Activating the p53 Pathway », Int. J. Mol. Sci., vol. 21, n° 23, p. 8949, nov. 2020, doi: 10.3390/ijms21238949.

[21] K. Ding et al., « RNA splicing factor USP39 promotes glioma progression by inducing TAZ mRNA maturation », Oncogene, vol. 38, n°37, p. 6414–6428, sept. 2019, doi: 10.1038/s41388-019-0888-1.

[22] J. Wu et al., « USP39 regulates DNA damage response and chemo-radiation resistance by deubiquitinating and stabilizing CHK2 », Cancer Lett., vol. 449, p. 114–124, mai 2019, doi: 10.1016/j.canlet.2019.02.015.

[23] Y. Peng et al., « USP39 Serves as a Deubiquitinase to Stabilize STAT1 and Sustains Type I IFN– Induced Antiviral Immunity », J. Immunol., vol. 205, n° 11, p. 3167–3178, déc. 2020, doi: 10.4049/jimmunol.1901384.

[24] Z. Zhang, W. Liu, T. Sun, J. Wang, M. Li, et C. Liu, « USP39 facilitates breast cancer cell proliferation through stabilization of FOXM », p. 22.

[25] Y. Xiao et al., « USP39-mediated deubiquitination of Cyclin B1 promotes tumor cell proliferation and glioma progression », Transl. Oncol., vol. 34, p. 101713, août 2023, doi: 10.1016/j.tranon.2023.101713.

[26] X. Li et al., « Deubiquitinase USP39 and E3 ligase TRIM26 balance the level of ZEB1 ubiquitination and thereby determine the progression of hepatocellular carcinoma », Cell Death Differ., vol. 28, n° 8, p. 2315–2332, août 2021, doi: 10.1038/s41418-021-00754-7.

[27] M.-A. Hamouda et al., « The small heat shock protein B8 (HSPB8) confers resistance to bortezomib by promoting autophagic removal of misfolded proteins in multiple myeloma cells ».

[28] W. E. Berndtson, « A simple, rapid and reliable method for selecting or assessing the number of replicates for animal experiments. », J. Anim. Sci., vol. 69, n° 1, p. 67, 1991, doi: 10.2527/1991.69167x.

[29] P. G. Richardson et al., « Bortezomib or High-Dose Dexamethasone for Relapsed Multiple Myeloma », N. Engl. J. Med., 2005.

[30] P. G. Richardson et al., « A Phase 2 Study of Bortezomib in Relapsed, Refractory Myeloma », N. Engl. J. Med., vol. 348, n° 26, p. 2609–2617, juin 2003, doi: 10.1056/NEJMoa030288.

[31] Y. Zhang, L. Xu, A. Li, et X. Han, « The roles of ZEB1 in tumorigenic progression and epigenetic modifications », Biomed. Pharmacother., vol. 110, p. 400–408, févr. 2019, doi: 10.1016/j.biopha.2018.11.112.

[32] A. Sacco et al., « Cancer Cell Dissemination and Homing to the Bone Marrow in a Zebrafish Model », Cancer Res., vol. 76, n° 2, p. 463–471, janv. 2016, doi: 10.1158/0008-5472.CAN-15-1926.

[33] A. M. Antao, A. Tyagi, K.-S. Kim, et S. Ramakrishna, « Advances in Deubiquitinating Enzyme Inhibition and Applications in Cancer Therapeutics », Cancers, vol. 12, n° 6, p. 1579, juin 2020, doi: 10.3390/cancers12061579.

[34] L. M. Doherty et al., « Integrating multi-omics data reveals function and therapeutic potential of deubiquitinating enzymes », eLife, vol. 11, p. e72879, juin 2022, doi: 10.7554/eLife.72879.

[35] N. Li et al., « Plasma cell myeloma with *RAS/BRAF* mutations is frequently associated with a complex karyotype, advanced stage disease, and poorer prognosis », Cancer Med., vol. 12, n° 13, p. 14293–14304, juill. 2023, doi: 10.1002/cam4.6103.

[36] J. M. Fraile et al., « USP39 Deubiquitinase Is Essential for KRAS Oncogene-driven Cancer », J. Biol. Chem., vol. 292, n° 10, p. 4164–4175, mars 2017, doi: 10.1074/jbc.M116.762757.

[37] H. Li et al., « Deubiquitylase USP12 induces pro-survival autophagy and bortezomib resistance in multiple myeloma by stabilizing HMGB1 », Oncogene, vol. 41, n°9, p. 1298–1308, févr. 2022, doi: 10.1038/s41388-021-02167-9.

[38] D. Chauhan et al., « A Small Molecule Inhibitor of Ubiquitin-Specific Protease-7 Induces Apoptosis in Multiple Myeloma Cells and Overcomes Bortezomib Resistance », Cancer Cell, vol. 22, n°3, p. 345–358, sept. 2012, doi: 10.1016/j.ccr.2012.08.007.

[39] Y. Song et al., « Blockade of deubiquitylating enzyme Rpn11 triggers apoptosis in multiple myeloma cells and overcomes bortezomib resistance », Oncogene, vol. 36, n° 40, p. 5631–5638, oct. 2017, doi: 10.1038/onc.2017.172.

[40] Y. Alsayed et al., « Mechanisms of regulation of CXCR4/SDF-1 (CXCL12)–dependent migration and homing in multiple myeloma », Blood, vol. 109, n° 7, p. 2708–2717, avr. 2007, doi: 10.1182/blood-2006-07-035857.

[41] G. Pagnucco, G. Cardinale, et F. Gervasi, « Targeting Multiple Myeloma Cells and Their Bone Marrow Microenvironment », Ann. N. Y. Acad. Sci., vol. 1028, n° 1, p. 390–399, déc. 2004, doi: 10.1196/annals.1322.047.

[42] M. Iwatsuki et al., « Epithelial–mesenchymal transition in cancer development and its clinical significance », Cancer Sci., vol. 101, n° 2, p. 293–299, févr. 2010, doi: 10.1111/j.1349-7006.2009.01419.x.

[43] D. Greaves et Y. Calle, « Epithelial Mesenchymal Transition (EMT) and Associated Invasive Adhesions in Solid and Haematological Tumours », Cells, vol. 11, n°4, p. 649, févr. 2022, doi: 10.3390/cells11040649.

[44] Y. Peng et al., « IGF-1 promotes multiple myeloma progression through PI3K/Akt-mediated epithelial-mesenchymal transition », Life Sci., vol. 249, p. 117503, mai 2020, doi: 10.1016/j.lfs.2020.117503.

[45] « Biological roles and prognostic values of the epithelial mesenchymal transition-mediating (1).pdf ».

[46] V. Stavropoulou et al., « MLL-AF9 Expression in Hematopoietic Stem Cells Drives a Highly Invasive AML Expressing EMT-Related Genes Linked to Poor Outcome », Cancer Cell, vol. 30, n° 1, p. 43–58, juill. 2016, doi: 10.1016/j.ccell.2016.05.011.

[47] I. M. Ghobrial, « Myeloma as a model for the process of metastasis: implications for therapy », Blood, vol. 120, n° 1, p. 20–30, juill. 2012, doi: 10.1182/blood-2012-01-379024.

[48] A. K. Azab et al., « Hypoxia promotes dissemination of multiple myeloma through acquisition of epithelial to mesenchymal transition-like features », Blood, vol. 119, n° 24, p. 5782–5794, juin 2012, doi: 10.1182/blood-2011-09-380410.

[49] C. M. Cheong et al., « Twist-1 is upregulated by NSD2 and contributes to tumour dissemination and an epithelial-mesenchymal transition-like gene expression signature in t(4;14)-positive multiple myeloma », Cancer Lett., vol. 475, p. 99–108, avr. 2020, doi: 10.1016/j.canlet.2020.01.040.

[50] Z. Zhou et al., « USP51 promotes deubiquitination and stabilization of ZEB », p. 12.

[51] L. Sun, J. Yu, J. Guinney, B. Qin, et F. A. Sinicrope, « USP10 Regulates ZEB1 Ubiquitination and Protein Stability to Inhibit ZEB1-Mediated Colorectal Cancer Metastasis », Mol. Cancer Res., vol. 21, n° 6, p. 578–590, juin 2023, doi: 10.1158/1541-7786.MCR-22-0552.

[52] K. Zeng et al., « USP22 upregulates ZEB1-mediated VEGFA transcription in hepatocellular carcinoma », Cell Death Dis., vol. 14, n° 3, p. 194, mars 2023, doi: 10.1038/s41419-023-05699-y.

[53] C. Song et al., « USP18 promotes tumor metastasis in esophageal squamous cell carcinomas via deubiquitinating ZEB1 », Exp. Cell Res., vol. 409, n° 1, p. 112884, déc. 2021, doi: 10.1016/j.yexcr.2021.112884.

[54] D. Ye, S. Wang, Y. Huang, X. Wang, et P. Chi, « USP43 directly regulates ZEB1 protein, mediating proliferation and metastasis of colorectal cancer », J. Cancer, vol. 12, n° 2, p. 404–416, 2021, doi: 10.7150/jca.48056.

